# HIPK2-and IKKβ-dependent phosphorylation stabilizes TAp63α during the oocyte DNA damage response

**DOI:** 10.64898/2026.04.17.719163

**Authors:** Anjali Kumari, Kiran Kumar P, Aradhana Mohanty, Abhilasha S, Lava Kumar S, Ajith Kumar E, Mohd Athar, Pravin Birajdar, Akshay Kumar, Silambresan Y, Sahina Sabnam, H.B.D. Prasada Rao

## Abstract

TAp63α, a member of the p53 family, serves as a central quality control factor in oocytes, safeguarding genomic integrity during the prolonged dictyate arrest of meiosis. In healthy oocytes, TAp63α is maintained in an inactive dimeric state; upon DNA damage, it undergoes phosphorylation-dependent tetramerization, enabling transcriptional activation of pathways that determine oocyte fate. While the upstream activation cascade of TAp63α has been well characterized, the mechanisms that regulate its stability during the DNA damage response remain incompletely understood. Here, we identify the kinases HIPK2 and IKKβ as key regulators of TAp63α stability. We show that TAp63α interacts with both kinases and is phosphorylated at distinct residues, T452 by HIPK2 and S4/S12 by IKKβ in vitro and in vivo, in addition to previously described CHK2 and CK1-mediated phosphorylation. Functionally, these phosphorylation events do not primarily contribute to activation, but instead stabilize TAp63α by limiting MDM4-dependent ubiquitination and subsequent proteasomal degradation. Mechanistically, our data support a model in which CHK2 and CK1 initiate TAp63α phosphorylation, while HIPK2 and IKKβ act in a complementary manner to maintain protein stability during genotoxic stress. Disruption of HIPK2 or IKKβ activity reduces TAp63α stability, whereas their inhibition in vivo attenuates oocyte loss following DNA damage, resulting in increased preservation of the follicle pool. Importantly, these effects are observed across multiple systems, including mouse models and ex vivo goat ovary cultures, supporting an evolutionarily conserved role for this regulatory axis. Together, our findings uncover a previously unidentified layer of TAp63α regulation, in which phosphorylation not only contributes to its activation but also enhances protein stability, thereby fine-tuning oocyte responses to DNA damage. Our results further indicate that HIPK2 and IKKβ-mediated phosphorylation modulates oocyte survival under genotoxic stress, highlighting this pathway as a potential target for strategies aimed at limiting oocyte loss.

## Introduction

The P53 family of proteins, TP53, TP63, and TP73, control cell cycle progression, DNA repair, apoptosis, and other cellular functions. TP63, in particular, is involved in the development of different tissues, including the skin, limbs, and organs (Chillemi et al., 2017; Gebel et al., 2017; Marcel et al., 2011). The TP63 gene is transcribed from two distinct promoters, resulting in either TAp63 isoforms with an N-terminal transactivation (TA) domain or ΔNp63 isoforms lacking the TA domain. Further, the 3′ ends of both TAp63 and ΔNp63 transcripts are alternately spliced to produce the various C-terminal isoforms known as α, β and γ(Fisher et al., 2020; Moll and Slade, 2004; Osterburg and Dötsch, 2022; Petitjean et al., 2007). The transactivating isoforms of TAp63 have an N-terminal transactivation domain (TAD) structurally identical to the transactivation domain of TP53, which enables them to control gene expression by binding to specific DNA regions and initiating transcription(Candi et al., 2014). There are three TA isoforms: TAp63α, TAp63β, and TAp63γ(Petitjean *et al*., 2007). Out of these three isoforms, TAp63α is expressed in the female germline(Suh et al., 2006). Specifically, TAp63α is expressed in mouse oocytes from primordial, primary, and early secondary follicles despite the fact that it is not required for follicle maturation, ovulation, or fertility(Luan et al., 2021). Recent research indicates that ionizing radiation causes DNA damage in primordial and primary oocytes, resulting in mortality(Gebel et al., 2020). Surprisingly, TAp63α -deficient primordial and primary oocytes survive, suggesting that TAp63α plays an important role in DNA damage response(Luan et al., 2022).

The DNA damage response (DDR) in oocytes is comparable to that of somatic cells (Carroll and Marangos, 2013). During prenatal oocyte DNA damage, ataxia telangiectasia mutant (ATM) and ataxia telangiectasia and Rad-3 related (ATR) proteins are activated and phosphorylate histone H2 (H2AX) at serine 139, allowing the other DDR proteins to assemble at DNA damage sites and activate CHK2(Collins and Jones, 2016). Further, CHK2 phosphorylates P53, causing transcriptional activation of downstream targets of cell cycle arrest, repair, or apoptosis genes. However, P53 is replaced by TAp63α after birth, specifically in primordial, primary, and secondary oocytes, to activate downstream targets transcriptionally(Emori et al., 2023; Rinaldi et al., 2020; Singh et al., 2023). Mechanistically, TAp63α will be in a closed dimeric state with very low DNA binding affinity in primordial, primary, and secondary oocytes. Soon after DNA damage, the CHK2-dependent phosphorylation at S582 primes and switches the TAp63α from a closed to an open dimeric state. Further, the CK1 executes the phosphorylation at remaining sites S585, S588, S591, and T594 of TAp63α to promote active tetramerization. The TAp63α tetramerization increases the affinity for DNA and transcriptional machinery(Tuppi et al., 2018). Previous findings imply that regulated DNA damage-dependent TAp63α tetramerization is essential in oocyte mortality(Deutsch et al., 2011; Tuppi *et al*., 2018). The resistance of TAp63 null mice oocytes to gamma radiation suggests that TAp63α is required to regulate oocyte death(Livera et al., 2008; Suh *et al*., 2006). In contrast, wild-type or P53 null oocytes died entirely, showing that TAp63α phosphorylation appeared to be a critical step in the response to DNA damage(Suh *et al*., 2006). In addition to CHK2 and CK1, prior research suggests that the tyrosine kinase c-Abl phosphorylates TAp63α at Tyr149, Tyr171, and Tyr289 in response to DNA damage(Gonfloni, 2010). Additionally, the Abl-kinase inhibitor imatinib effectively prevents the activation of TAp63α and protects mouse oocytes from cisplatin-induced damage(Gonfloni et al., 2009). This observation suggests that, besides CHK2 and CK1, other kinases may play a role in TAp63α phosphorylation. Previous research on different p63 isoforms has demonstrated that IKKβ-mediated phosphorylation of TAp63γ enhances its stability by inhibiting its ubiquitylation and subsequent degradation(Liao et al., 2013; MacPartlin et al., 2008). In contrast, HIPK2, associated with apoptosis and drug response, phosphorylates and targets the ΔNp63α isoform, an anti-apoptotic variant for degradation, thereby increasing chemosensitivity(Lazzari et al., 2011). Despite these findings, the specific roles of HIPK2 and IKKβ in regulating TAp63α remain to be elucidated. This study reveals that TAp63α is phosphorylated by HIPK2 and IKKβ kinases, in addition to CHK2 and CK1, contributing to its stabilization and the maintenance of follicular pool.

## Results

### TAp63α interacts with HIPK2 and IKKβ in ovaries and the H1299 cell line

To investigate how TAp63α is regulated in the female germline, we aimed to define its protein interactome. Given the logistical and ethical challenges associated with using large cohorts of mice, we employed goat ovaries, as the TAp63α antibody demonstrates effective cross-reactivity with goat TAp63α. This choice was supported by the robust cross-reactivity of the TAp63α antibody with the goat protein. Immunohistochemical analysis confirmed that TAp63α is specifically localized in primordial, primary, and secondary follicles (Fig. S1A and B). Based on this, we selected ovarian samples enriched for TAp63α-positive follicles to maximize the yield and relevance of downstream interaction studies.

TAp63α immunoprecipitation was then performed and validated by western blotting, followed by mass spectrometry to identify potential interacting partners (Fig. 1A, B, and Fig. S1C). The analysis identified known regulators of TAp63α, including CHK2 and CK1, supporting the reliability of the approach. In addition, HIPK2 and IKKβ were identified as potential novel interacting partners (Fig. 1C and Fig. S1C), suggesting previously unrecognized regulatory inputs into the TAp63α pathway. However, further validation of these interactions in goat tissue was limited by the lack of antibodies that reliably cross-react with goat HIPK2 and IKKβ.

**Figure 1.**
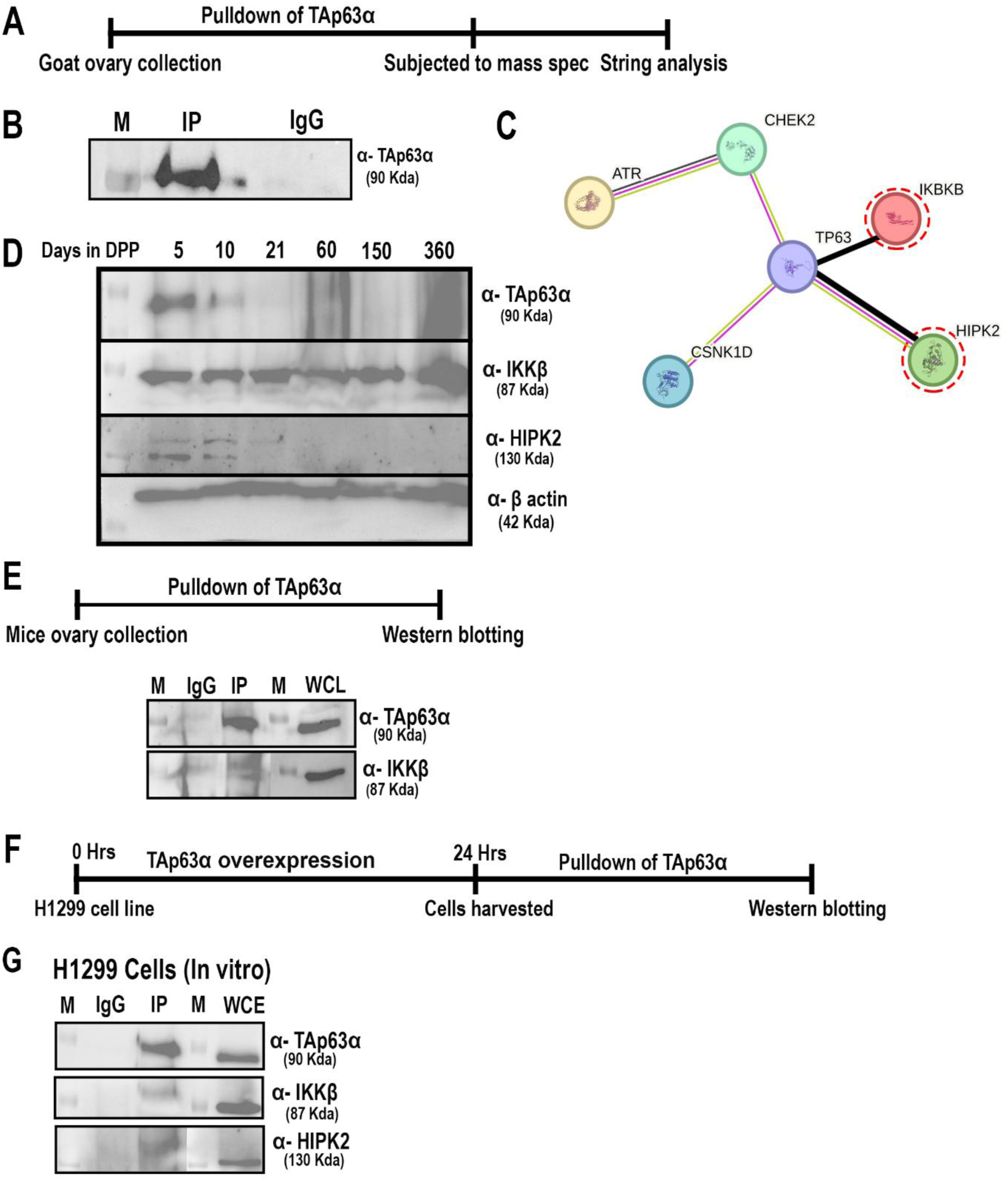
Identification and validation of HIPK2 and IKKβ as TAp63α-interacting partners. (A) Schematic overview of the experimental workflow used to identify TAp63α-interacting proteins from ovarian tissue. (B) Western blot analysis confirming successful immunoprecipitation of endogenous TAp63α from goat ovary lysates. (C) STRING network analysis of proteins identified by mass spectrometry, highlighting predicted interactions between TAp63α and the kinases HIPK2 and IKKβ. (D) Western blot analysis of ovarian lysates from mice of different ages showing that HIPK2 and TAp63α protein levels decrease with age, whereas IKKβ levels remain relatively constant. Two pairs of ovaries were pooled per sample. (E) Co-immunoprecipitation of endogenous TAp63α from mouse ovaries (eight pairs pooled per sample), followed by immunoblot detection of TAp63α and associated IKKβ. (F) Schematic representation of the experimental workflow used for interaction validation in a cell-based system. (G) Co-immunoprecipitation analysis in H1299 cells transiently expressing TAp63α, confirming interaction with HIPK2 and IKKβ. Whole-cell extracts and immunoprecipitates were probed with antibodies against TAp63α, HIPK2, and IKKβ. Due to differences in protein abundance and detection sensitivity, blots were processed and exposed separately to ensure optimal signal detection.

To overcome this limitation, we returned to mouse ovaries and investigated the expression of HIPK2 and IKKβ across different developmental stages, ranging from 5 days postpartum (dpp) to 1 year of age. Western blot analysis revealed that TAp63α was prominently expressed between 5 dpp and 21 dpp, whereas IKKβ was present throughout the observed time points. In contrast, HIPK2 exhibited a similar temporal expression pattern to TAp63α, being present from 5 to 21 dpp. These findings suggest that both TAp63α and HIPK2 are primarily involved in the early stages of folliculogenesis (Fig.1D). In mice, TAp63α is predominantly localized within primordial and primary follicles (Fig. S1B). As mice age, the number of TAp63α-positive follicles declines, which may explain the absence of detectable TAp63α protein beyond 21dpp. This reduction is likely attributable to the decreased expression of TAp63α at more advanced follicular stages, coinciding with the overall reduction in primordial and primary follicle populations over time.

Further co-immunoprecipitation experiments in mouse ovaries confirmed the interaction between TAp63α and IKKβ, but HIPK2 was not detected in the TAp63α co-IP (Fig.1E). To overcome this limitation, we transiently overexpressed TAp63α in H1229 cells carrying a mutated P53 and performed immunoprecipitation for TAp63α (Fig.1F). Subsequent co-IP with HIPK2 and IKKβ from the eluates revealed that both proteins were present, confirming that TAp63α interacts with HIPK2 and IKKβ under these conditions (Fig.1G, Fig. S1D and E).

### HIPK2 and IKKβ-dependent phosphorylation stabilizes TAp63α during the DNA damage response

To examine the functional relevance of TAp63α phosphorylation by HIPK2 and IKKβ, we transiently expressed TAp63α in cultured cells and subjected them to DNA damage in the presence or absence of kinase inhibitors targeting CHK2, CK1, HIPK2, or IKKβ. Cells were harvested 30 hours post-treatment, and protein lysates were analyzed by western blotting (Fig. 2A).

**Figure 2.**
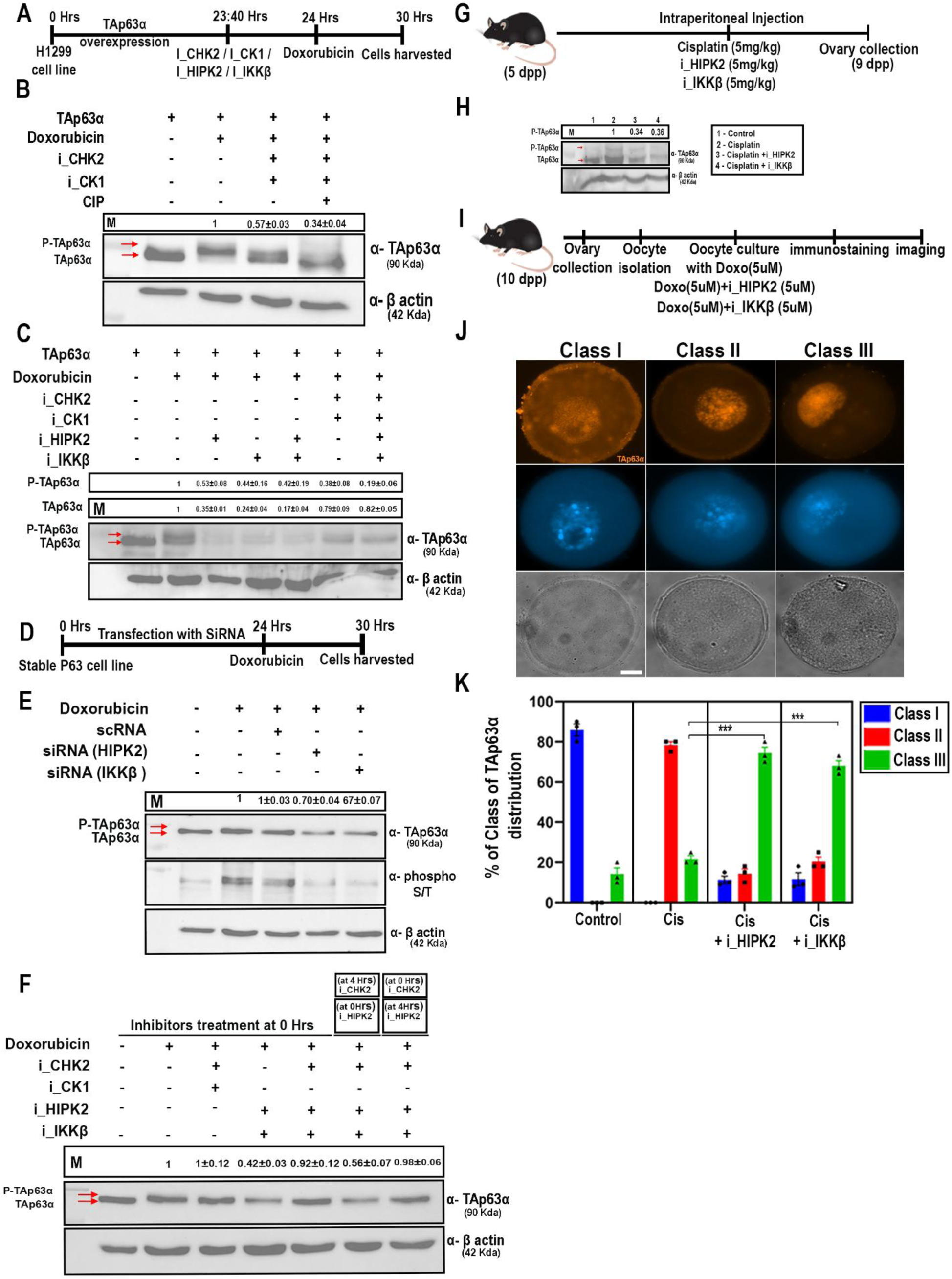
HIPK2- and IKKβ-dependent phosphorylation stabilizes TAp63α in a CHK2-dependent manner. (A) Schematic overview of the experimental approach used to assess kinase-dependent regulation of TAp63α following DNA damage. (B) Western blot analysis showing a phosphorylation-dependent mobility shift of TAp63α upon doxorubicin treatment. This shift is reduced by CHK2 and CK1 inhibition and is further diminished following calf intestinal phosphatase (CIP) treatment, confirming phosphorylation-dependent modification. (C) Inhibition of HIPK2 or IKKβ, individually or in combination, reduces TAp63α phosphorylation and is associated with decreased protein stability. In contrast, inhibition of CHK2 or CK1 reduces phosphorylation without affecting total TAp63α levels. (D) Schematic representation of the siRNA-mediated knockdown strategy in stable TAp63α-expressing H1299 cells. (E) Knockdown of HIPK2 or IKKβ reduces the phosphorylation-associated mobility shift of TAp63α compared to control and scrambled siRNA conditions. Immunoblotting with phospho-serine/threonine antibodies further confirms a reduction in overall phosphorylation levels upon kinase depletion. (F) Western blot analysis of TAp63α following inhibition of CHK2, HIPK2, and IKKβ under DNA damage conditions. CHK2 inhibition reduces the phosphorylation-associated mobility shift of TAp63α. Inhibition of HIPK2 and IKKβ decreases total TAp63α protein stability. Co-inhibition of CHK2 with HIPK2 and IKKβ restores TAp63α protein levels when applied prior to or concurrently, but not when CHK2 inhibition is applied after HIPK2 and IKKβ inhibition. (G) Schematic representation of the in vivo experimental workflow. (H) Western blot analysis of mouse ovarian lysates showing that cisplatin induces a phosphorylation-associated mobility shift of TAp63α. Co-treatment with HIPK2 or IKKβ inhibitors reduces both phosphorylation and total TAp63α levels. Five pairs of ovaries were pooled per sample. (I) Schematic overview of the experimental approach used for oocyte analysis. (J–K) Representative images showing distinct patterns of TAp63α nuclear distribution in oocytes, along with their quantification. Treatment with HIPK2 or IKKβ inhibitors under DNA damage conditions results in a shift toward reduced nuclear signal intensity. Statistical significance for Class III follicles: doxorubicin + HIPK2 inhibitor (P = 0.0001) and doxorubicin + IKKβ inhibitor (P = 0.0001) compared to doxorubicin alone. Data are presented as mean ± SD; unpaired t-test.

Upon treatment with the genotoxic agent doxorubicin, TAp63α exhibited a clear electrophoretic mobility shift, indicative of phosphorylation (Fig. 2B). Inhibition of CHK2 or CK1 under DNA damage conditions reduced this mobility shift, confirming their established roles as upstream kinases in TAp63α activation. However, treatment with calf intestinal phosphatase (CIP) further diminished the phosphorylated forms, suggesting either incomplete inhibition of CHK2/CK1 or the involvement of additional kinases (Fig. 2B). Increasing inhibitor concentrations was not feasible due to cytotoxicity. Notably, inhibition of HIPK2 or IKKβ, either individually or in combination, not only reduced TAp63α phosphorylation but also led to a pronounced decrease in total TAp63α protein levels (Fig. 2C). In contrast, inhibition of CHK2 or CK1 reduced phosphorylation without affecting protein stability (Fig. 2C). These observations indicate that, unlike CHK2 and CK1, HIPK2 and IKKβ play a critical role in maintaining TAp63α stability during the DNA damage response. To exclude potential off-target effects of chemical inhibitors, we validated these findings using siRNA-mediated knockdown in stable TAp63α-expressing H1299 cells (Fig. 2D). Silencing of HIPK2 or IKKβ reduced the phosphorylation-associated mobility shift compared to control and scrambled siRNA-treated cells (Fig. 2E). Furthermore, probing with anti-phospho-serine/threonine antibodies confirmed a reduction in overall phosphorylation levels upon HIPK2 or IKKβ depletion, consistent with the inhibitor-based results (Fig. 2E).

Previous studies have established CHK2 as the priming kinase and CK1 as the executioner kinase in TAp63α activation. We therefore asked whether HIPK2- and IKKβ-mediated phosphorylation depends on this upstream signaling cascade. Under DNA damage conditions, inhibition of CHK2 reduced TAp63α phosphorylation, as expected. Importantly, combined inhibition of CHK2 with HIPK2 or IKKβ further decreases phosphorylation levels (Fig. 2F), suggesting that multiple kinases contribute to the phosphorylation profile of TAp63α. Interestingly, the reduction in TAp63α protein stability observed upon HIPK2 or IKKβ inhibition was rescued when CHK2 was simultaneously inhibited (Fig. 2C, F). This indicates that CHK2 activity is required for the destabilization phenotype observed in the absence of HIPK2/IKKβ, pointing to an interdependent regulatory relationship.

To further dissect the sequence of kinase action, we performed sequential inhibitor treatments. When CHK2 was inhibited prior to or together with HIPK2/IKKβ inhibition, TAp63α stability was preserved despite reduced phosphorylation. In contrast, inhibition of HIPK2 and IKKβ prior to CHK2 inhibition failed to rescue protein stability (Fig. 2F). These findings suggest that CHK2-dependent priming phosphorylation is required for subsequent HIPK2/IKKβ-mediated stabilization of TAp63α. Together, these results support a model in which CHK2 and CK1 initiate TAp63α phosphorylation, while HIPK2 and IKKβ act to stabilize the phosphorylated protein during genotoxic stress.

To assess the physiological relevance of these findings in vivo, we treated 5-day postpartum (5 dpp) mice with cisplatin, either alone or in combination with HIPK2 or IKKβ inhibitors. Ovaries were collected at day 9, a stage at which TAp63α is readily detectable (Fig. 2G). Western blot analysis revealed that cisplatin induced a mobility shift consistent with TAp63α phosphorylation, whereas co-treatment with HIPK2 or IKKβ inhibitors reduced both phosphorylation and total protein levels (Fig. 2H).

We next examined TAp63α localization in oocytes by immunostaining. In cisplatin-treated ovaries, TAp63α showed strong pan-nuclear localization. However, inhibition of HIPK2 or IKKβ resulted in a shift toward weaker and more discontinuous nuclear staining patterns. Quantitative analysis revealed an increase in oocytes with weak TAp63α signal and a corresponding decrease in strongly stained nuclei (Fig. S2A, B and C), supporting a role for these kinases in maintaining TAp63α stability in vivo. Further, to directly assess early follicle responses, oocytes were isolated and subjected to in vitro DNA damage followed by treatment with HIPK2 or IKKβ inhibitors. Immunofluorescence analysis revealed altered TAp63α localization patterns, with a significant increase in condensed or diminished nuclear signals upon kinase inhibition (Fig. 2I–K). These changes further support that HIPK2 and IKKβ-mediated phosphorylation is essential for preserving TAp63α stability during genotoxic stress.

### TAp63α is phosphorylated at T452 by HIPK2 and at S4 and S12 by IKKβ

To validate the phosphorylation of TAp63α by HIPK2 and IKKβ, recombinant TAp63α was purified and subjected to in vitro phosphorylation assays using CHK2, CK1, HIPK2, IKKβ, or combinations thereof, along with radio-labelled ATP (Fig.3A, Fig.S3A and B). The resulting radiogram revealed that CHK2 robustly phosphorylates TAp63α (Fig.3B). CK1 alone showed minimal phosphorylation activity, whereas co-incubation of CHK2 and CK1 produced phosphorylation patterns similar to CHK2 alone, suggesting that CHK2 acts as the initiator kinase, with CK1 functioning as an executioner kinase (Fig.3B).

**Figure 3.**
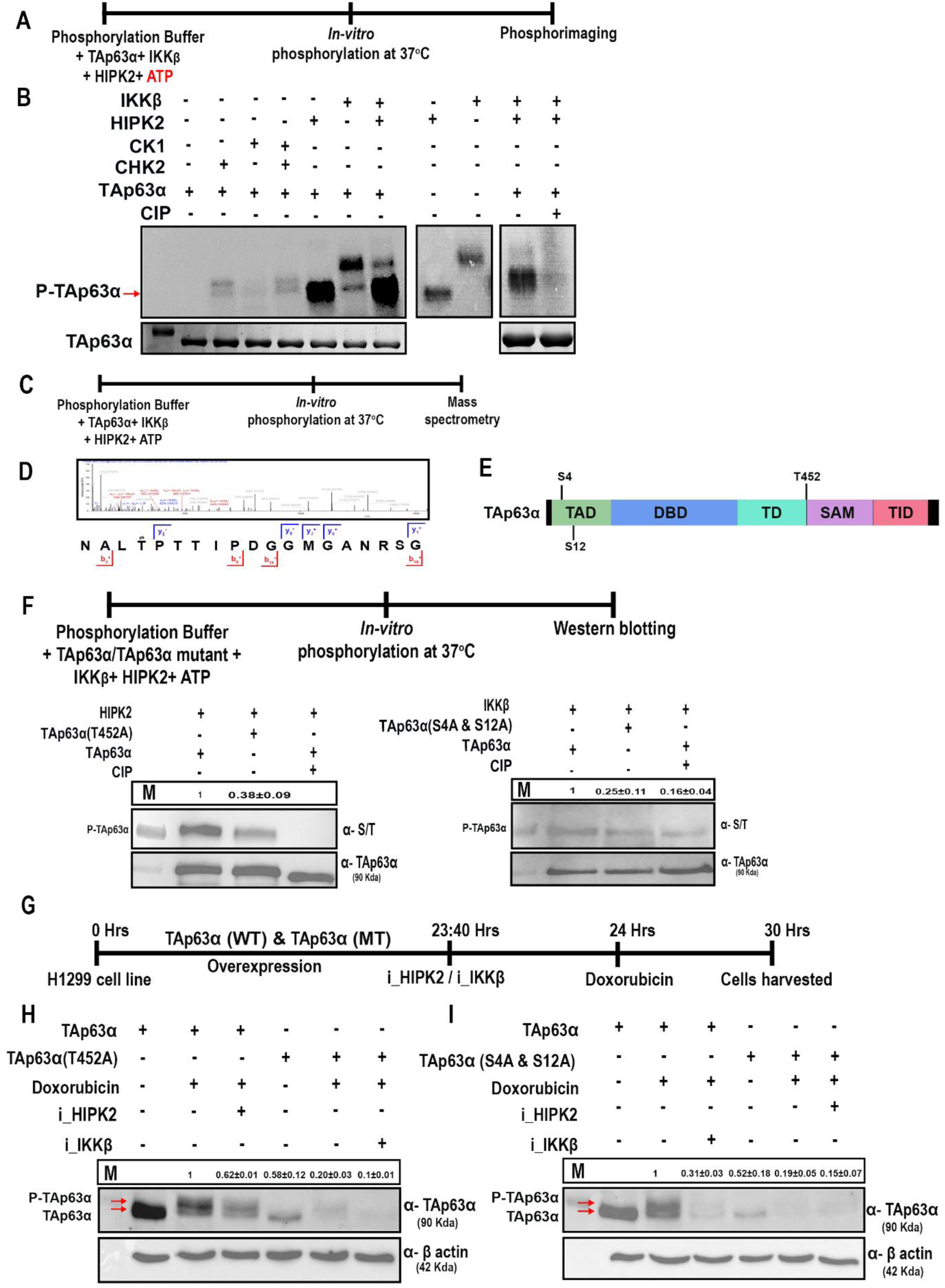
HIPK2 and IKKβ phosphorylate TAp63α in vitro at T 452 and S4 and S12. (A) Schematic representation of the experimental approach used to investigate TAp63α in vitro phosphorylation. (B) Autoradiogram demonstrating the phosphorylation of TAp63α by HIPK2 and IKKβ. Chk2, CK1, HIPK2, and IKKβ alone serve as positive controls. The addition of calf intestinal phosphatase (CIP) to the in vitro phosphorylation assay completely abolishes TAp63α phosphorylation. (C) Schematic depiction of the experimental workflow for mass spectrometry. (D) Mass spectrometry analysis identifying phosphorylation sites on TAp63α targeted by HIPK2 and IKKβ. (E) Schematic representation of TAp63α phosphorylation sites modified by HIPK2 and IKKβ. (F) Western blot analysis shows that T452A and S4A/S12A mutations reduce TAp63α phosphorylation levels. (G) Diagram illustrating the experimental methodology. (H) Western blot analysis indicates that the T452A mutation destabilizes TAp63α. (I) Western blot analysis demonstrates that the S4A and S12A mutations destabilize TAp63α. *unpaired* t*-test,* Data are presented as mean ± SD.

When HIPK2 or IKKβ were tested individually, both kinases phosphorylated TAp63α (Fig.3B). Co-incubation of HIPK2 and IKKβ resulted in an enhanced phosphorylation signal, indicating a synergistic effect (Fig.3B). CIP significantly reduced the phosphorylation, confirming that HIPK2 and IKKβ are responsible for TAp63α phosphorylation (Fig.3B).

Additionally, we performed auto-phosphorylation assays for HIPK2 and IKKβ to differentiate between their self-phosphorylation and their ability to phosphorylate TAp63α (Fig.3B). This analysis confirmed that the observed phosphorylation is dependent on HIPK2 and IKKβ activity rather than auto-phosphorylation.

We conducted an in vitro phosphorylation assay using cold ATP to identify specific amino acid residues in TAp63α targeted by HIPK2 and IKKβ (Fig.3C). Mass spectrometry analysis revealed that HIPK2 phosphorylates TAp63α at threonine 452 (T452) (Fig.3D, E and Fig.S3C). However, we did not detect any phosphorylation sites for IKKβ in vitro. Previous studies suggest that IKKβ phosphorylates serine 4 (S4) and serine 12 (S12) in the TAp63γ isoform (Fig.3E). To assess the functional significance of these phosphorylation sites, we generated recombinant TAp63α mutants in which key residues were substituted with alanine (T452A, S4A, and S12A) and subjected them to in vitro phosphorylation assays (Fig.3F). Western blot analysis confirmed that HIPK2 efficiently phosphorylates wild-type TAp63α, but phosphorylation was markedly reduced in the T452A mutant, indicating that T452 is a critical site for HIPK2-mediated phosphorylation (Fig.3F). Similarly, phosphorylation was significantly diminished in the S4A and S12A mutants in the presence of IKKβ, confirming that these residues are essential for IKKβ-mediated phosphorylation (Fig.3F).

We further investigated the role of specific phosphorylation sites by transiently transfecting H1229 cells with wild-type or mutant (T452A, S4A, S12A) TAp63α constructs, followed by treatment with the DNA-damaging agent doxorubicin to assess phosphorylation and protein stability (Fig.3G). Western blot analysis demonstrated a phosphorylation-induced mobility shift in wild-type TAp63α upon DNA damage (Fig.3H). However, inhibition of HIPK2 significantly reduced this shift and decreased protein stability. The T452A mutant exhibited reduced basal expression levels, even without DNA damage (Fig.3H). Upon DNA damage or combined DNA damage and HIPK2 inhibition, the T452A mutant displayed further destabilization, indicating that HIPK2-mediated phosphorylation of T452 is critical for maintaining TAp63α stability under genotoxic stress (Fig.3H). Similarly, the S4A and S12A mutants showed lower basal expression levels (Fig.3I). These mutants were further destabilized upon DNA damage with IKKβ inhibition, suggesting that IKKβ-mediated phosphorylation of S4 and S12 is crucial for stabilizing TAp63α during genotoxic stress (Fig.3I).

### HIPK2- and IKKβ-mediated phosphorylation stabilizes TAp63α by limiting MDM4-dependent ubiquitination

To investigate the mechanism by which HIPK2- and IKKβ-mediated phosphorylation regulates TAp63α stability, we first tested whether this modification protects TAp63α from proteasomal degradation. TAp63α was transiently overexpressed in H1299 cells and subjected to DNA damage in the presence or absence of HIPK2 or IKKβ inhibitors, with or without the proteasome inhibitor MG132 (Fig. 4A). Western blot analysis revealed that DNA damage induced a phosphorylation-associated mobility shift of TAp63α. Inhibition of HIPK2 or IKKβ reduced both phosphorylation and total TAp63α protein levels. Importantly, co-treatment with MG132 restored TAp63α protein levels despite kinase inhibition (Fig. 4B), indicating that loss of HIPK2/IKKβ activity may promote proteasome-dependent degradation of TAp63α.

**Figure 4.**
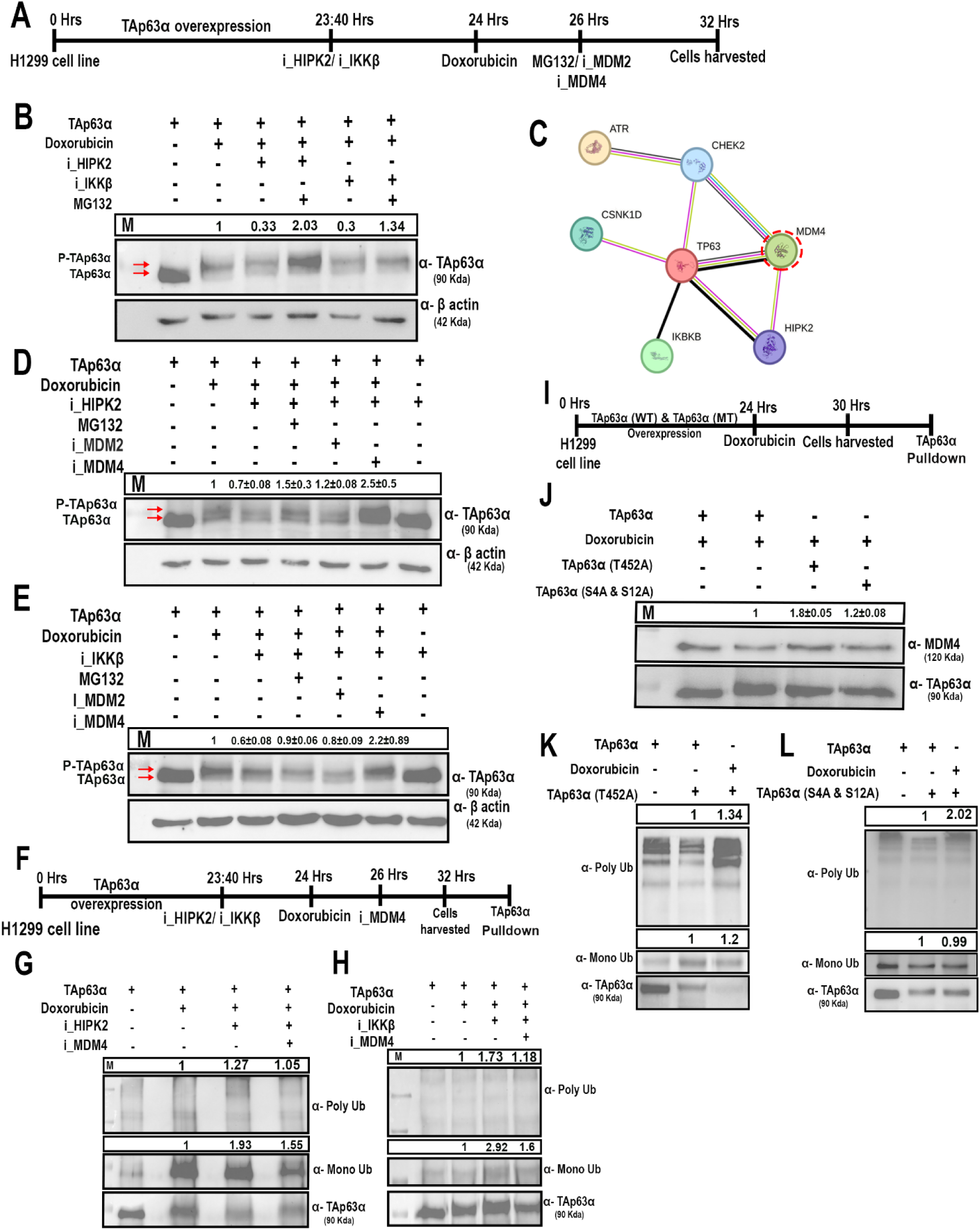
HIPK2 and IKKβ-dependent phosphorylation controls TAp63α ubiquitination and stability. (A) Schematic overview of the experimental workflow. (B) Western blot analysis showing that inhibition of HIPK2 and IKKβ reduces TAp63α phosphorylation and total protein levels. Treatment with the proteasome inhibitor MG132 restores TAp63α protein levels. (C) Protein interaction network analysis identifying MDM4 as a potential TAp63α-interacting partner. (D–E) Western blot analysis of TAp63α levels following inhibition of MDM4 or MDM2 in the presence of HIPK2 and IKKβ inhibitors. MDM4 inhibition restores TAp63α protein levels, whereas MDM2 inhibition shows a partial effect. (F) Schematic representation of the experimental design for ubiquitination assays. (G–H) Western blot of ubiquitination assays showing increased mono- and poly-ubiquitination of TAp63α upon inhibition of HIPK2 and IKKβ. Co-inhibition of MDM4 reduces TAp63α ubiquitination levels. (I) Schematic overview of the experimental approach for interaction analysis. (J) Co-immunoprecipitation analysis showing interaction between TAp63α and MDM4 under basal and DNA damage conditions. Reduced interaction is observed following DNA damage in wild-type TAp63α, whereas non-phosphorylatable mutants retain interaction with MDM4. (K–L) Western blot of ubiquitination assays comparing wild-type and phosphorylation-deficient TAp63α mutants, showing increased ubiquitination of the mutant proteins. Data are presented as mean ± SD; statistical significance determined by unpaired t-test.

Because proteasomal degradation is typically mediated by ubiquitination, we next sought to identify the E3 ubiquitin ligase responsible for targeting TAp63α. Reanalysis of our goat TAp63α interactome identified MDM4 as a potential interacting partner (Fig. 4C; Fig. S4). To determine the functional role of MDM4 in regulating TAp63α stability, we performed inhibitor-based experiments targeting HIPK2, IKKβ, MDM4, and MDM2 under DNA damage conditions (Fig. 4D). Consistent with earlier observations, inhibition of HIPK2 or IKKβ reduced both phosphorylation and protein levels of TAp63α. Co-inhibition of MDM4 in this context resulted in a robust restoration of TAp63α levels, whereas inhibition of MDM2 had minimal effect (Fig. 4D, E). These findings indicate that MDM4, rather than MDM2, is the primary E3 ligase mediating TAp63α degradation, and that HIPK2/IKKβ-dependent phosphorylation counteracts this process.

We next directly assessed TAp63α ubiquitination under these conditions. Immunoprecipitation of TAp63α followed by ubiquitin detection revealed that DNA damage induces both mono-and poly-ubiquitination of TAp63α (Fig. 4F). Notably, inhibition of HIPK2 or IKKβ further increased both mono- and poly-ubiquitinated forms (Fig. 4G, H), consistent with enhanced targeting for degradation. In contrast, co-treatment with MDM4 inhibitors markedly reduced ubiquitination levels, even in the presence of HIPK2 or IKKβ inhibition. These results demonstrate that HIPK2- and IKKβ-mediated phosphorylation restrains MDM4-dependent ubiquitination of TAp63α during genotoxic stress.

To further dissect how phosphorylation influences the interaction between TAp63α and MDM4, we compared wild-type TAp63α with non-phosphorylatable mutants (T452A and S4A/S12A). Under basal conditions, wild-type TAp63α interacted with MDM4, consistent with the proteomic data. However, upon DNA damage, this interaction was markedly reduced (Fig. 4I, J), suggesting that phosphorylation weakens MDM4 binding. In contrast, the non-phosphorylatable mutants retained or even showed enhanced interaction with MDM4 under DNA damage conditions (Fig. 4J). Consistent with this, ubiquitination assays revealed that the phosphorylation-deficient mutants exhibited increased ubiquitination compared to wild-type TAp63α (Fig. 4K, L). These findings indicate that phosphorylation of TAp63α at HIPK2 and IKKβ-targeted residues disrupts its interaction with MDM4, thereby limiting ubiquitination and promoting protein stability. Together, these results demonstrate that HIPK2 and IKKβ-mediated phosphorylation stabilizes TAp63α by preventing its recognition and ubiquitination by MDM4, thereby protecting it from proteasomal degradation during the DNA damage response.

### HIPK2 and IKKβ modulate oocyte response to DNA damage

Since HIPK2 and IKKβ-dependent phosphorylation of TAp63α is crucial for its stabilization, and TAp63α phosphorylation plays a key role in the oocyte DNA damage response, we aimed to investigate the functional significance of these kinases in the follicle dynamics and oocyte survival. To understand their roles, we administered pharmacological inhibitors targeting either HIPK2 or IKKβ, individually and in combination, to pre-pubertal and post-pubertal mice (Fig.5A and D).

**Figure 5.**
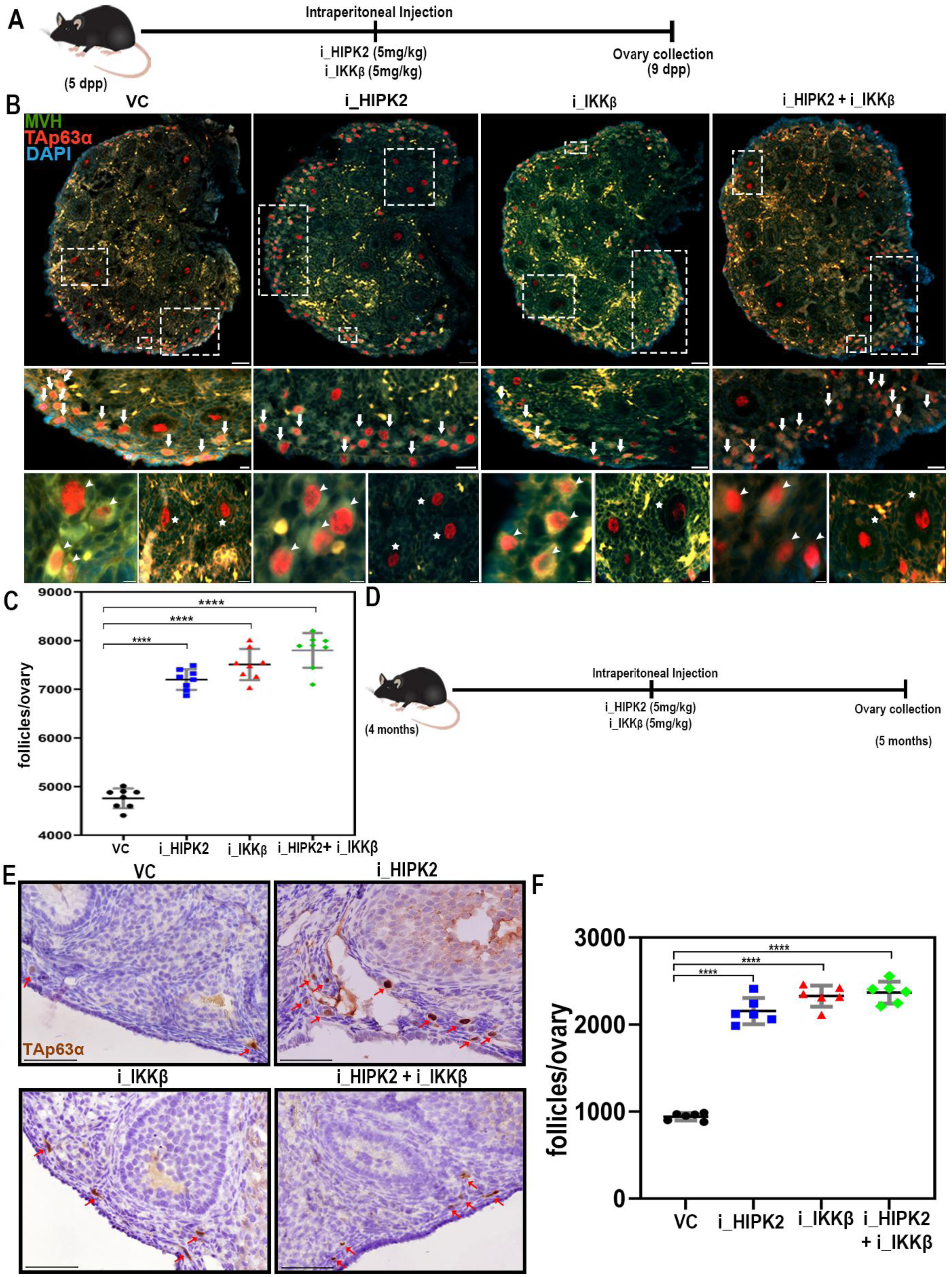
HIPK2 and IKKβ are essential for follicle pool maintenance. (A) Schematic representation of the experimental workflow. (B) Representative images of ovaries from 9-day postpartum (dpp) C57BL/6 mice injected with five doses of different treatments: DMSO (control), HIPK2 inhibitor, IKKβ inhibitor, or a combination of both. Ovaries were immunostained for MVH (green, marking germ cells), TAp63α (red, indicating guardian of germline), and DNA (blue, nuclear staining). (C) Quantification of the follicles. Statistical analysis shows significant protection of follicles in all treatment groups compared to the control (HIPK2 inhibitor: *P ≤ 0.0001*, IKKβ inhibitor: *P ≤ 0.0001*, combination treatment: *P ≤ 0.0001*). (D) Schematic illustration of the experimental methodology. (E) Representative images of ovaries from 5-month-old C57BL/6 mice injected twice weekly with DMSO (control), HIPK2 inhibitor, IKKβ inhibitor, or their combination. Ovaries were immunostained for TAp63α (brown) with hematoxylin counterstaining. (F) Quantification of the follicles in 5-month-old mice. Statistical analysis reveals significant protection in all treatment groups compared to the control (*P ≤ 0.0001* for HIPK2 inhibitor, IKKβ inhibitor, and combination treatment). *unpaired* t*-test,* Data are presented as mean ± SD. Scale bars for 9dpp mice ovary section: 100μm and for 5month mice ovary section: 50μm.

In pre-pubertal mice (5dpp), we administered 5 mg/kg body weight of the inhibitors daily up to 9 days. Following treatment, the animals were sacrificed, and the ovaries were collected, fixed, sectioned, and immunostained with the germline marker MVH and the guardian of the germline, TAp63α (Fig.5A and B). Total follicle numbers were quantified. Results showed that inhibition of HIPK2 resulted in 1.5-fold more follicles compared to controls, while inhibition of IKKβ led to a 1.6-fold increase in follicle count, suggesting that inhibition of either kinase preserves the ovarian follicles (Fig.5C and Fig. S5A). Interestingly, simultaneous inhibition of both kinases did not further increase follicle numbers, indicating that HIPK2 and IKKβ may function within the same regulatory pathway in this context (Fig.5C).

Similarly, in post-pubertal mice (90 dpp), inhibitors were administered twice weekly for 30 days (Fig.5D). Ovaries were collected at 120 days, fixed, and subjected to immunohistochemistry with TAp63α (Fig.5E). Quantification of follicle numbers revealed that inhibition of HIPK2 increased follicle numbers by 2.3-fold compared to controls, while inhibition of IKKβ resulted in a 2.5-fold increase (Fig.5F and Fig. S5B). As observed in pre-pubertal mice, co-inhibition of HIPK2 and IKKβ yielded similar results to single kinase inhibition, suggesting that both kinases operate through a shared pathway in regulating follicle preservation during aging (Fig.5F).

To evaluate the impact of HIPK2 and IKKβ inhibition on oocyte, embryo quality and reproductive potential, we administered specific inhibitors from 5 dpp to day 9 dpp, followed by a two-month maturation period. After this interval, one cohort underwent oocyte and embryo retrieval and in vitro maturation (IVM) assays, while a separate cohort was assessed for fertility through mating with untreated males (Fig.S6A and B). Oocyte maturation analysis, tracking progression from the germinal vesicle (GV) to metaphase II-like (MII), revealed no significant differences between treated and control groups (Fig.S6C and D). The embryo maturation analysis shows no significant difference (Fig. S6E). Likewise, fertility assessments demonstrated no detrimental effects on mating success or litter size (Fig. S6G). Notably, anti-Müllerian hormone (AMH) levels, a key indicator of ovarian reserve, were elevated in treated animals compared to controls (Fig.S6I). To further assess the long-term consequences of early kinase inhibition, mice were treated transiently during the pre-pubertal window (5–9 dpp) and allowed to age without further intervention. Remarkably, analysis at 60 dpp revealed a sustained ∼2.2-fold increase in follicle numbers in inhibitor-treated groups (Fig. S5C-E), demonstrating that early modulation of this pathway has lasting protective effects on the follicles. These findings indicate that the transient inhibition of HIPK2 and IKKβ preserves follicles without impairing oocyte competence or reproductive outcomes.

### Inhibition of HIPK2 and IKKβ protects oocytes from genotoxic stress and preserves the follicle pool

To determine the physiological role of HIPK2 and IKKβ in regulating oocyte survival under genotoxic stress, we examined the impact of their inhibition on follicle survival using both pre-pubertal and post-pubertal mouse models. Five-day postpartum (5 dpp) mice were administered cisplatin, either alone or in combination with inhibitors of HIPK2, IKKβ, or both, on alternate days. Ovaries were collected at 9 dpp and analyzed by immunostaining for MVH and TAp63α (Fig. 6A, B).

**Figure 6.**
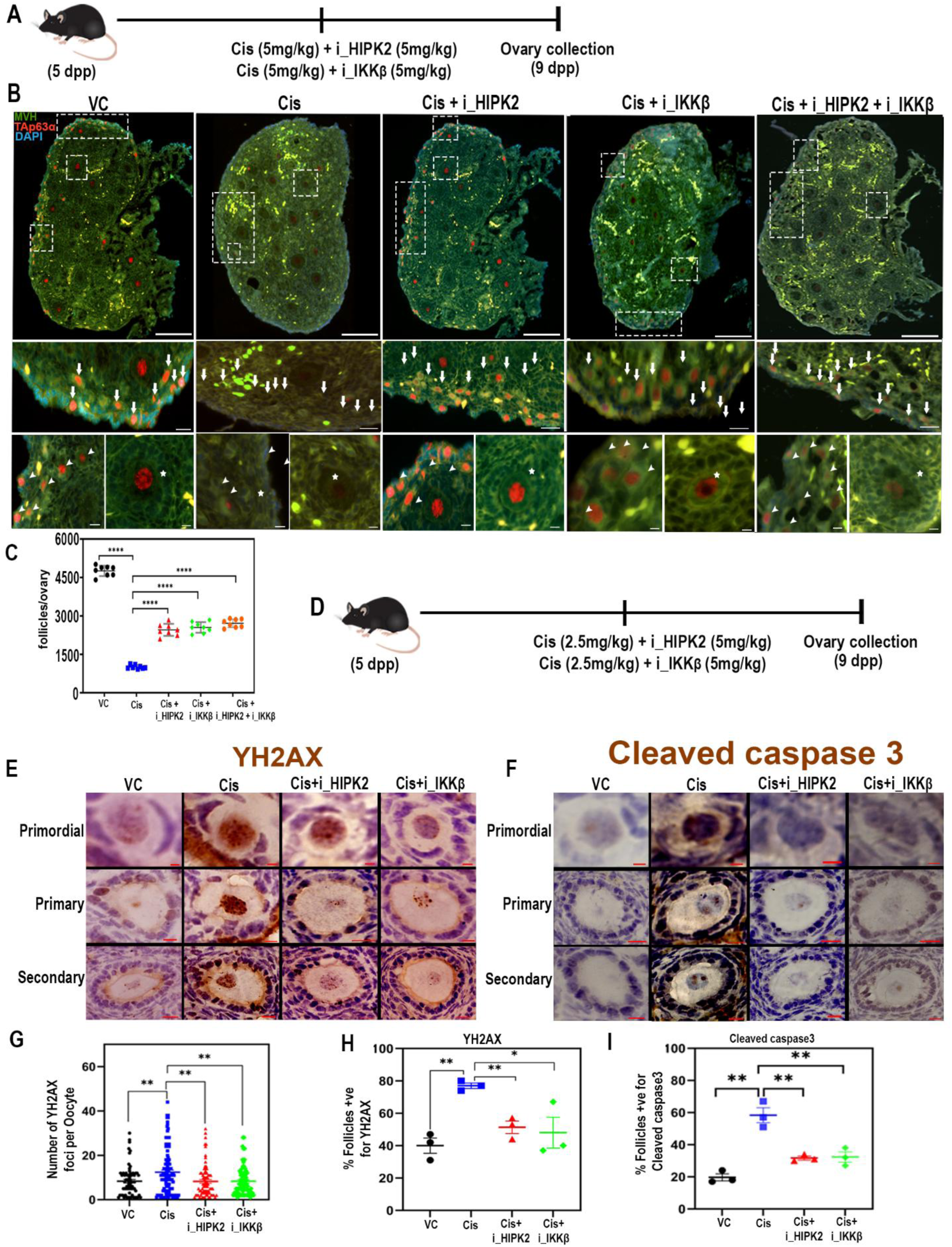
Inhibition of HIPK2 and IKKβ preserves the follicle pool during chemotherapy. (A) Schematic representation of the experimental design. (B) Representative images of ovarian sections from 9-day-postpartum (dpp) C57BL/6 mice treated with Vehicle control (VC), Cisplatin, Cisplatin + HIPK2 inhibitor, Cisplatin + IKKβ inhibitor, and Cisplatin + HIPK2 & IKKβ inhibitors. Ovaries were immunostained for MVH (green, marking germ cells), TAp63α (red, marking DNA damage response), and DNA (blue, nuclear staining). (C) Quantification of the follicles from panel (B). Statistical analysis shows significant differences in follicle counts for Vehicle Control (*P = 0.0001*), Cisplatin + HIPK2 inhibitor (*P = 0.0001*), Cisplatin + IKKβ inhibitor (*P = 0.0013*), and Cisplatin + combined inhibitors (*P = 0.0001*) compared to the only Cisplatin group. (D) Schematic representation of the experimental design. Representative images of primordial, primary and secondary follicles from ovarian sections of 9-day-postpartum (dpp) C57BL/6 mice treated with Vehicle control, Cisplatin, Cisplatin + HIPK2 inhibitor, Cisplatin + IKKβ inhibitor. Ovaries were immunostained for gH2AX (brown) (E) and for cleaved caspase 3 (F) (brown) with hematoxylin counterstaining. Quantification of the (G) number of gH2AX foci per Oocyte. Statistical significance in comparison to cisplatin group: Vehicle control (*P = 0.0034*), Cisplatin + HIPK2 inhibitor (*P = 0.0023*), Cisplatin + IKKβ inhibitor (*P = 0.0020*) (H) % of follicles positive for gH2AX. Statistical significance in comparison to cisplatin group: Vehicle control (*P = 0.0018*), Cisplatin + HIPK2 inhibitor (*P = 0.0037*), Cisplatin + IKKβ inhibitor (*P = 0.04*) (I) % of follicles positive for cleaved caspase 3. Statistical significance in comparison to cisplatin group: Vehicle control (*P = 0.0017*), Cisplatin + HIPK2 inhibitor (*P = 0.0052*), Cisplatin + IKKβ inhibitor (*P = 0.010*). *unpaired* t*-test,* Data are presented as mean ± SD. Scale bars for 9dpp mice ovary section: 100μm, Primordial and primary follicles:10μm, Secondary follicles:5μm.

Quantitative analysis revealed that cisplatin treatment caused a marked depletion of the follicle pool, with approximately a 4.7fold reduction in follicle numbers compared to controls, reflecting extensive oocyte loss due to DNA damage (Fig. 6C). In contrast, co-treatment with either HIPK2 or IKKβ inhibitors significantly mitigated this loss, resulting in only a 2.5fold reduction in follicle numbers. Combined inhibition of both kinases did not confer additional protection beyond single treatments, suggesting that HIPK2 and IKKβ function within the same regulatory pathway to control oocyte fate during genotoxic stress (Fig. 6C, Fig. S7A).

A similar protective effect was observed in post-pubertal mice. Sixty-day-old mice treated with cisplatin exhibited a comparable three-fold reduction in follicle numbers, whereas co-treatment with HIPK2 or IKKβ inhibitors preserved the follicle pool. (Fig. S7B-E). Again, dual inhibition did not further enhance protection, reinforcing the idea that these kinases act in a common pathway.

To understand the underlying cellular mechanisms, we assessed DNA damage and apoptosis in ovarian sections by immunostaining for γH2AX and cleaved caspase-3 (CC3) (Fig. 6D–F and Fig. S8). As expected, cisplatin treatment significantly increased both the number of γH2AX foci and the proportion of γH2AX-positive follicles, indicating elevated DNA damage in early-stage follicles (Fig. 6E, G, H). Notably, inhibition of HIPK2 or IKKβ reduced both the intensity and frequency of γH2AX-positive oocytes compared to cisplatin alone, suggesting an attenuation of the DNA damage response or improved oocyte survival. Consistent with this, apoptosis analysis revealed that approximately 60% of follicles were positive for cleaved caspase-3 following cisplatin treatment. In contrast, co-treatment with HIPK2 or IKKβ inhibitors led to an approximately 1.5-fold reduction in apoptotic follicles (Fig. 6F, I), indicating that inhibition of these kinases suppresses oocyte apoptosis under genotoxic stress. At the molecular level, gene expression analysis further supported this protective effect. Pro-apoptotic genes such as *PUMA* and *NOXA* were downregulated, whereas anti-apoptotic genes including *Bcl2l1* and *Mcl1* were upregulated in ovaries co-treated with kinase inhibitors and cisplatin compared to cisplatin alone (Fig. S9A, B). These changes are consistent with reduced TAp63α activity due to its destabilization, leading to a dampened transcriptional response to DNA damage. Together, these findings demonstrate that inhibition of HIPK2 and IKKβ attenuates DNA damage signaling and apoptosis in oocytes, thereby preserving the follicle pool under genotoxic stress.

### Evolutionarily conserved HIPK2- and IKKβ-dependent control of follicle preservation under genotoxic stress

To investigate the evolutionarily conserved mechanisms of HIPK2 and IKKβ in follicle pool maintenance, we utilized goat ovaries at the late fetal stage (30 cm crown-rump length), a developmental period characterized by abundant early follicle formation enriched with TAp63α expression. Ovaries were subjected to an air-liquid interface culture system and treated with or without HIPK2 and IKKβ inhibitors for 48 hours. Following the culture, ovaries were harvested, fixed, and immunostained for markers of germ cells (MVH) and TAp63α (Fig. 7A). Control ovarian sections revealed a significant loss of follicles, indicative of natural follicular attrition during culture (Fig. 7B and C). In contrast, ovaries treated with HIPK2 and IKKβ inhibitors displayed a 2-fold and 4-fold increase in follicle survival, suggesting that inhibition of these kinases plays a protective role in maintaining the follicle pool (Fig. 7B and C). Gene expression analysis shown downregulation of Pro-apoptotic gene bbc3 whereas upregulation of anti-apoptotic gene *Mcl1* in goat oocytes co-treated with kinase inhibitors and doxorubicin compared to doxorubicin alone (Fig. S9C, D). These findings demonstrate that HIPK2 and IKKβ are critical regulators of follicular attrition, and their inhibition preserves the follicle pool, a mechanism that appears to be evolutionarily conserved across species.

**Figure 7.**
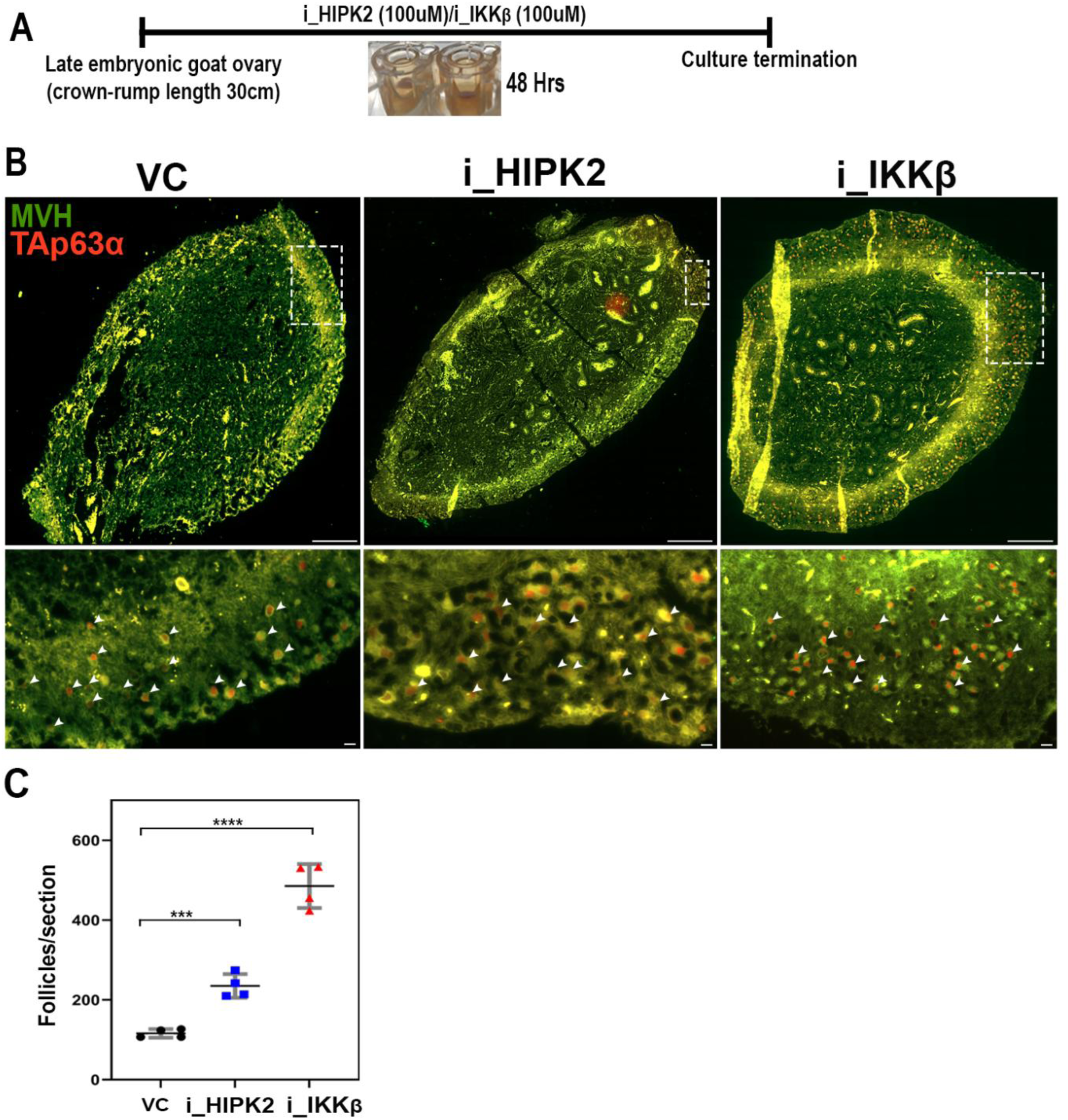
Evolutionary conservation of HIPK2- and IKKβ-dependent follicle pool maintenance. (A) Schematic representation of the ex vivo ovarian culture method used for goat ovaries. (B) Representative images of goat ovarian sections immunostained for MVH (green) and TAp63α (red) under different treatment conditions: DMSO (control), HIPK2 inhibitor, and IKKβ inhibitor. Enlarged views of the respective ovarian sections are shown below. (C) Quantification of the follciles from panel B. Statistical significance in comparison to control: HIPK2 inhibitor \*\*\*\**P ≤ 0.0012*; IKKβ inhibitor \*\*\*\**P ≤ 0.0001* (*unpaired t-test*). Data are presented as mean ± SD. Scale bars: 200μm.

## Discussion

Our study shows that the phosphorylation-dependent stabilization of TAp63α is critical in preserving the follicle pool (model). TAp63α, a member of the p53 family of tumour suppressors, is essential for DNA damage-induced apoptosis in oocytes, ensuring the elimination of genetically compromised cells(Gebel *et al*., 2020). In the present study, we identified two kinases, HIPK2 and IKKβ, as pivotal regulators of TAp63α stability. These kinases play distinct roles in maintaining oocyte integrity and protecting the follicle pool under both basal and genotoxic stress conditions (model). Our findings expand on the established role of TAp63α in DNA damage response pathways by showing that phosphorylation by HIPK2 and IKKβ enhances its stability, thereby protecting the follicle quality.

### HIPK2 and IKKβ mediated phosphorylation of p63 isoforms: Cell specific functions and mechanisms

HIPK2 and IKKβ, well-known for their roles in DNA damage response pathways, modulate the activity and stability of various p63 isoforms via phosphorylation(Lazzari *et al*., 2011; Liao *et al*., 2013; MacPartlin *et al*., 2008). The p63 protein exists in multiple isoforms, including TAp63 and ΔNp63, each with unique functions in regulating development, apoptosis, and cellular responses to stress(Osterburg and Dötsch, 2022). TAp63α, the isoform predominantly expressed in female germ cells, plays a critical role in safeguarding oocytes by initiating apoptosis in response to DNA damage(Luan *et al*., 2022; Tuppi *et al*., 2018). Our study demonstrated that HIPK2 phosphorylates TAp63α at threonine 452, while IKKβ targets serines 4 and 12. These phosphorylation events stabilize TAp63α by inhibiting MDM4-mediated ubiquitination, typically marking proteins for proteasomal degradation. Consequently, stabilized TAp63α may form active tetramers with high DNA-binding affinity, leading to the transcriptional activation of DNA repair or pro-apoptotic genes to protect or eliminate damaged oocytes (model). This mechanism maintains genomic integrity in the female germline, particularly under genotoxic conditions such as chemotherapy.

In contrast to the role of TAp63α in oocytes, IKKβ stabilizes the TAp63γ isoform in somatic cells, enhancing its transcriptional activity and inducing cell cycle arrest and apoptosis under stress conditions, suggesting a conserved mechanism where IKKβ phosphorylation fine tunes the function of distinct TAp63 isoforms depending on the cellular environment(Liao *et al*., 2013; MacPartlin *et al*., 2008). Meanwhile, HIPK2 phosphorylates the ΔNp63α isoform, which lacks the N-terminal transactivation domain and functions as a dominant-negative regulator of the p53 family(Lazzari *et al*., 2011). HIPK2-mediated phosphorylation of ΔNp63α promotes its degradation via ubiquitination, thereby lifting its inhibitory effect on pro-apoptotic pathways, contrary to the stabilizing role of TAp63α in oocytes(Lazzari *et al*., 2011). Thus, the phosphorylation of TAp63α by HIPK2 and IKKβ highlights a complex regulatory network that governs p63 isoform activity. In oocytes, the stabilization of TAp63α is crucial for triggering DNA repair or apoptosis in response to DNA damage, whereas, in somatic cells, HIPK2 and IKKβ modulate other isoforms to influence cell fate decisions under stress.

### A multi-kinase architecture safeguards TAp63α stability and calibrates follicle pool maintenance

Under physiological conditions, TAp63α is maintained in an inactive dimeric state within oocytes, poised to respond to genotoxic insults. Upon DNA damage, a well-defined phosphorylation cascade initiated by CHK2 and amplified by CK1 drives its rapid conversion into an active tetramer, thereby triggering transcriptional programs that eliminate damaged oocytes (Gebel et al., 2020). While this activation module is essential for enforcing genomic integrity, it does not fully explain how TAp63α protein levels are sustained during this response. Our study identifies HIPK2 and IKKβ as critical components of a previously unrecognized regulatory layer that governs TAp63α stability.

Mechanistically, we show that HIPK2 and IKKβ phosphorylate TAp63α at distinct residues (T452, S4, and S12), which collectively act to antagonize MDM4-mediated ubiquitination and subsequent proteasomal degradation. This stabilization ensures that once activated, TAp63α is maintained at levels sufficient to execute its function. Importantly, this reveals that TAp63α regulation is not limited to activation kinetics but extends to active control of protein turnover. In vivo evidence further supports this model, as inhibition of HIPK2 or IKKβ disrupts TAp63α stability and significantly alters follicle dynamics under genotoxic stress, underscoring the physiological relevance of this pathway.

These findings support a two-tiered regulatory framework for TAp63α function. The CHK2–CK1 axis constitutes an upstream activation module that governs the switch-like transition of TAp63α into its active tetrameric state. In contrast, HIPK2 and IKKβ form a downstream stabilization module that determines the persistence of the activated protein by shielding it from ubiquitin-mediated degradation. This division of labor highlights that activation and stability are mechanistically separable yet functionally interdependent processes.

The presence of both HIPK2 and IKKβ in this stabilization module is not simply redundant but reflects a design principle of robustness and signal integration. Oocytes are uniquely sensitive cells that must interpret varying degrees and types of DNA damage over extended periods. The involvement of multiple kinases likely ensures that TAp63α stabilization is tightly coupled to the intensity and duration of genotoxic stress. Such redundancy provides resilience, allowing partial compensation if one pathway is compromised, while also enabling fine-tuning of the response.

Functionally, this architecture introduces a critical buffering system that calibrates the threshold for oocyte elimination. While CHK2–CK1 rapidly activates TAp63α in response to DNA lesions, HIPK2 and IKKβ determine whether this activation is sustained or transient. In conditions where HIPK2/IKKβ activity is limited, TAp63α becomes unstable and is degraded, thereby dampening its pro-apoptotic output. This controlled destabilization prevents excessive oocyte loss under moderate or transient stress, preserving the follicle pool. Conversely, under severe or persistent damage, sustained stabilization of TAp63α ensures efficient elimination of compromised oocytes. Taken together, our findings redefine TAp63α regulation as a dynamic balance between activation and stability, orchestrated by a multi-kinase network. Rather than acting redundantly, HIPK2 and IKKβ function as critical modulators that integrate stress signals and fine-tune oocyte fate decisions, thereby safeguarding both genomic integrity and reproductive longevity.

### Implications for fertility preservation

The finite nature of the oocytes and its vulnerability to genotoxic stress pose significant challenges to fertility preservation, especially in women undergoing cancer treatments(Mahajan, 2015). The ability to pharmacologically target pathways that stabilize TAp63α presents a potential strategy for mitigating oocyte loss during chemotherapy(Singh *et al*., 2023). Our findings suggest that inhibitors of HIPK2 and IKKβ could be explored as therapeutic agents to prevent premature ovarian failure and extend fertility in cancer patients. Furthermore, our experiments using goat ovaries revealed the conservation of this mechanism across species, underscoring the potential translational impact of targeting HIPK2 and IKKβ in fertility preservation strategies. The stabilization of TAp63α through HIPK2 and IKKβ-mediated phosphorylation offers a promising avenue for protecting the oocytes and extending the reproductive lifespan under both physiological and pathological conditions.

## Materials and methods

### 1.1 Expression and purification of recombinant TAp63α

The TAp63α codon-optimized for expression in *E. coli* was cloned into the pET30a (+) vector and purchased from GenScript. *E. coli* BL21 (DE3) cultures expressing TAp63α were induced with IPTG, harvested by centrifugation at 4°C, and resuspended in lysis buffer (50 mM Tris-HCl, pH 8.0, 300 mM NaCl, 0.1% SDS, PMSF, protease inhibitor, and 10 mM imidazole). Cells were lysed by sonication, and the lysate was centrifuged at 14,000 rpm for 30 minutes at 4°C. The supernatant was collected and initially purified using Ni-NTA chromatography, with protein elution performed using 200–500 mM imidazole. The eluted protein was then dialyzed against 50 mM Tris and 100 mM NaCl, flash-frozen, and stored at -80°C until use in *in vitro* reactions.

### 1.2 Site-Directed Mutagenesis

Site-directed mutagenesis was performed by designing mutant primers for all target sites using the Oligo Calc primer design tool. Serine residues at positions 4 and 12, as well as the threonine residue at position 452 of TAp63α, were mutated to alanine using mutant primers in a PCR reaction. The methylated template plasmid DNA was digested with DpnI at 37°C for 3 hours and subsequently transformed into DH10β cells. The plasmid was then isolated and sequenced to confirm the mutations.

### 1.3 Phosphorylation Assays

A total of 1 μg of purified TAp63α was incubated with γ-³²P-ATP (2 μCi) in the presence of HIPK2 (∼0.2 μg) (Cat. No. PV5275, Thermo) and/or IKKβ (∼0.2 μg) (Cat. No. PV3836, Thermo) in a 20 μl reaction buffer containing 50 mM Tris-HCl (pH 8.0), 0.1 mM EDTA, 2 mM DTT, 10 mM MgCl₂, 10 mM β-glycerophosphate, and 10 mM sodium fluoride. The reaction mixture was incubated at 30°C for 1 hour, after which the reaction was terminated by adding 5× Laemmli SDS-PAGE sample buffer. The samples were then subjected to electrophoresis on a 10% SDS-PAGE gel. Following electrophoresis, the gel was placed in contact with a phosphor screen to capture radioactive emissions. The phosphor screen was subsequently scanned using a phosphorimager to visualize the results.

### 2.1 Cell Culture

The human lung adenocarcinoma cell line H1299 (ATCC - CRL-5803) was obtained from ATCC and cultured in RPMI 1640 medium (Cat no. 11875085, Gibco) supplemented with 10% fetal bovine serum (FBS) (Cat no. 10270106, Gibco) and 1× penicillin-streptomycin (Cat no. P4333, Sigma) at 37°C in a humidified incubator with 5% CO₂. Transfection was performed using the p63 plasmid (Cat no. 27008, Addgene) following the Lipofectamine™ 3000 Transfection Reagent protocol (Cat no. L3000008, Thermo). After 24 hours of transfection, the cells were centrifuged at 15,000 rpm for 10 minutes at 4°C to collect the pellet. To evaluate the effects of various inhibitors, the following compounds were introduced 23 hours and 40 minutes post-TAp63α transfection: Chk2 inhibitor (i_Chk2) (10 μM; BML-277, Cat no. 17552, Cayman), CK1 inhibitor (i_ck1) (1 μM; PF-670462, Cat no. 14588, Cayman), HIPK2 inhibitor (i_HIPK2) (10 μM; Protein kinase inhibitors 1, Cat no. HY-U00439A, MedChem), and IKKβ inhibitor (i_IKKβ) (10 μM; IKK16, Cat no. HY-13687, MedChem). After 20 minutes, DNA damage was induced using doxorubicin (10 μM; Cat no. 15007, Cayman). Subsequently, 2 hours after doxorubicin treatment, additional inhibitors were introduced, including the MDM4 inhibitor (i_MDM4) (30 μM; NSC-207895, Cat no. HY-14714, MedChem), MDM2 inhibitor (i_MDM2) (30 μM; Nutlin-3, Cat no. 10009816, Cayman), and proteasome inhibitor (30 μM; MG-132, Cat no. M3244, TCI). The culture was then terminated 32 hours post-transfection. Finally, the cells were centrifuged at 15,000 rpm for 10 minutes at 4°C to collect the pellet.

### 2.2 Stable TAp63α cell line generation

HEK293T packaging cells were plated 24 hours prior to transfection in a 10 cm culture dish with DMEM and 10% FBS, allowing them to reach 70–80% confluency. HEK293T cells were co-transfected with the lentiviral transfer plasmid pPRK01-p63-HFP-FLAG, the packaging plasmid psPAX2 (cat no.-12260, Addgene), and VSV-G (cat no.-14888, Addgene) using PEI at a DNA:PEI ratio of 1:3, and the mixture was incubated at room temperature for 15 minutes. The transfection solution was added dropwise to the HEK293T cells and kept overnight at 37°C in a humidified incubator with 5% CO₂. After 10 hours, the medium was replaced with fresh complete RPMI. Viral supernatants were harvested 48 hours after transfection, cleared by low-speed centrifugation to eliminate cellular debris, and filtered through a 0.45 µm syringe filter. The filtered lentiviral supernatant was either utilized immediately for transduction or stored at−80°C.

To generate stable cell lines, H1299 cells were plated in 6-well dishes at 40–50% confluency. The media was then exchanged for fresh complete RPMI media containing polybrene (3 µL per 1.5 mL medium). An equal volume of lentiviral supernatant (1.5 mL) was subsequently added to the cells, which were incubated for 24 hours. After the infection period, the viral medium was replaced with fresh complete RPMI media, allowing the cells to recover for another 24 hours. Around 48 hours post-infection, when the cells reached 80–90% confluency, the cells were trypsinized and replated in fresh complete RPMI media supplemented with blasticidin for antibiotic selection. The selection process continued until all non-transduced control cells were eliminated, and the resistant cells were stably expanded. GFP-positive cells were then further enriched using fluorescence-activated cell sorting. Stable TAp63α-expressing H1299 cells were subsequently expanded and cryopreserved.

Further, H1299 cells stably expressing TAp63α were prepared. Once the cells were 70-80% confluent, siRNA transfection was performed following the Lipofectamine™ 3000 Transfection Reagent protocol (Cat no. L3000008, Thermo). After 24 hours of transfection, DNA damage was induced using doxorubicin (10 μM; Cat no. 15007, Cayman) and the cells were harvested at 30^th^ hours, centrifuged at 15,000 rpm for 10 minutes at 4°C to collect the pellet.

For the mass spectrometry, H1299 cells stably expressing TAp63α with 70-80% confluency was treated with HIPK2 inhibitor (i_HIPK2) and IKKβ inhibitor (i_IKKβ) followed by doxorubicin treatment at 20^th^ min. cells were harvested after 6 hours of Doxo treatment and processed for mass spectrometry.

To further dissect the sequence of kinase action, we performed sequential inhibitor treatments. Once the H1299 cells stably expressing TAp63α were 70-80% confluent, at 0^th^ hour in one condition cells were treated with Chk2 inhibitor (i_Chk2), HIPK2 inhibitor (i_HIPK2) and IKKβ inhibitor (i_IKKβ) together, in 2^nd^ condition cells were treated first with HIPK2 inhibitor (i_HIPK2) and IKKβ inhibitor (i_IKKβ) together at 0^th^ hour and then with Chk2 inhibitor (i_Chk2) at 4^th^ hour & in 3^rd^ condition cells were treated first with only Chk2 inhibitor (i_Chk2) at 0^th^ hour and later HIPK2 inhibitor (i_HIPK2) and IKKβ inhibitor (i_IKKβ) together at 4^th^ hour. In all the condition DNA damage was introduced using doxorubicin at 20^th^ min. Inhibitor concentrations were used similar to as previously mentioned and the cells were harvested at 8^th^ hours, centrifuged at 15,000 rpm for 10 minutes at 4°C to collect the pellet.

## 3. CIP treatment

To dephosphorylate proteins using Calf Intestinal Alkaline Phosphatase (CIP), in vitro phosphorylated samples and cell lysates from doxorubicin-treated cells were prepared. The CIP enzyme (Cat. No. 2250B) was then added at a concentration of 5 units per microgram of protein. The reaction mixture was incubated at 37°C for 1 hour. To terminate the reaction, Laemmli buffer was added, followed by boiling at 100°C for 5 minutes. The samples were then loaded onto an SDS-PAGE gel for protein detection.

## 4. Co-immunoprecipitation

Goat ovaries, mouse ovaries, and cell pellets were lysed in a lysis buffer containing 25 mM Tris-HCl (pH 8.0), 150 mM NaCl, 1% Triton X-100, 1% Tween-20, 1% NP-40, PMSF, and a protease inhibitor. The lysates were then subjected to sonication and centrifuged at 12,000 rpm for 10 minutes at 4°C. Protein A/G beads (Cat. No. 88803, Thermo) were pre-washed with lysis buffer and incubated with anti-TAp63α antibody (Cat. No. CM 163B, Biocare) for 4 hours. Following antibody incubation, the beads were washed and subsequently incubated with the lysates for 16 hours. After incubation, the beads were washed and then resuspended in 1X Laemmli buffer, followed by incubation at 65°C for 30 minutes. The supernatant was collected, boiled at 100°C for 5 minutes, and loaded onto SDS-PAGE to analyze the protein of interest. Co-immunoprecipitated proteins were detected using western blot analysis with monoclonal antibodies against TAp63α (Cat. No. CM 163B, Biocare), HIPK2 (Cat. No. 5091, CST), IKKβ (Cat. No. 8943, CST), MDM4 (Cat. No. MO445, Sigma) and Ubiquitin (Cat. No. PAA164Mi01,Cloudclone).

To further check how phosphorylation and mutation influences the interaction between TAp63α and MDM4, we overexpressed wild type and mutant (T452A, S4A &S12A) TAp63α. Post 24 hours of transfection DNA damage was introduced using Doxorubicin and cells were harvested at 30^th^ hour. Cells were lysed and incubated with recombinant protein MDM4. Further pulldown of TAp63α was done from the cell lysate incubated with MDM4 as previously mentioned. Co-immunoprecipitated proteins were detected using western blot analysis with monoclonal antibodies against TAp63α (Cat. No. CM 163B, Biocare) and MDM4 (Cat. No. MO445, Sigma).

### 5.1 MS sample preparation

Goat ovaries were lysed in a lysis buffer, and TAp63α was pulled down following as described above. The elution was performed using 0.1 M glycine at pH 2.7, and the pH was subsequently adjusted by adding Tris-HCl at pH 9. The samples were then treated with 200 mM dithiothreitol (DTT) at 57°C for 1 hour to reduce disulfide bonds. Following this, alkylation was carried out by incubating the samples in the dark at room temperature with 200 mM iodoacetamide for 1 hour. Protein digestion was performed at 37°C for 16 hours using trypsin/LysC (Cat. No. V5073, Promega). The trypsin-digested samples were then treated with formic acid and subsequently purified using a C18 spin column (Cat. No. 89870, Thermo) according to the manufacturer’s instructions. After vacuum drying, the purified samples were reconstituted in 0.3% formic acid. A final concentration of 1 μg of peptide sample was injected into the mass spectrometer for analysis. Each experiment was conducted using two to three separate biological replicates.

### 5.2 Mass spectroscopy data analysis

Peptides and proteins were identified using Proteome Discoverer v2.5 software (Thermo Fisher Scientific, San José, CA, USA). The search parameters included dynamic modifications for oxidation at methionine residues and phosphorylation at serine, threonine, and tyrosine residues, as well as a static modification for carbamidomethylation at cysteine residues. The maximum number of missed cleavages was set to 2, with peptide length constraints defined by a maximum of 144 amino acids and a minimum of 6 amino acids as we previously published (Lava Kumar et al., 2024).

### 6.1 Animal experiments

The mice were maintained under a 12-hour light-dark cycle with ad libitum access to food and water, in accordance with the guidelines set by the National Institute of Animal Biotechnology Ethics Committee. All experiments were performed using C57BL/6 mice. The study commenced with 5-day postpartum (dpp) C57BL/6 mice, which were divided into four groups: vehicle control (VC), HIPK2 inhibitor (I_HIPK2; protein kinase inhibitor 1, Cat No. HY-U00439A, MedChem), IKKβ inhibitor (I_IKKβ; IKK16, Cat No. HY-13687, MedChem), and a combination of both inhibitors (I_HIPK2 and I_IKKβ). Each group consisted of 3–5 animals, and the mice received five intraperitoneal injections (5 mg/kg body weight) of either the inhibitors or the vehicle control, administered once daily in the morning from 5 dpp to 9 dpp. Ovaries were collected on the evening of 9 dpp and subsequently fixed. For the aging experiment similar regime was followed till 9 dpp and mice were left till 2month. After 2month animals were sacrificed and subsequently fixed.

Similarly, in post-pubertal mice (4 months old), the inhibitors were administered twice weekly for 30 days, following the same four-group experimental design. Ovaries were collected at 5 months and fixed.

For the DNA damage and rescue experiment, 5 dpp pups were injected with 5 mg/kg body weight of I_HIPK2, I_IKKβ, or a combination of both on the 5th, 7th, and 9th days, while cisplatin (Cat No. HY-17394, 5 mg/kg body weight) was administered on the 6th and 8th days in the morning. On the evening of 9 dpp, the pups were sacrificed, and their ovaries were collected and fixed. Similarly, in post-pubertal mice (60 dpp), inhibitors were administered on three alternate days, while cisplatin was administered on two days. Ovaries were collected at 65 dpp and fixed.

For the fertility assessment, 5 dpp mice received five doses (5 mg/kg body weight) of the inhibitors (VC, I_HIPK2, I_IKKβ, and the combination of I_HIPK2 and I_IKKβ), after which they were aged until 2 months and subsequently subjected to mating trials.

### 6.2 Oocyte Culture and Immunostaining

10 dpp FVB mice were used to collect the ovary, and ovarian puncture was performed using a 24-gauge needle in M2 medium (Cat. No. M7167, Sigma) to retrieve oocytes. The oocytes were isolated, washed, and subsequently kept in M2 medium with HIPK2 inhibitor (i_HIPK2-5 μM) and IKKβ inhibitor (i_IKKβ-5μM) along with control at 0^th^ hour, within a CO₂ incubator maintained at 37°C with 90–95% humidity. After 20 min DNA damage was introduced with doxorubicin (5 μM), following 6 hours of treatment oocytes were washed three times with media and fixed with 4% PFA for 30 min. After fixation oocytes were washed once with 1XTBST-0.05% Tween 20 (0.05% TBST), permeabilisation was done with 1XTBST-0.1% Triton-X-100 for 15 min, followed by washing twice with 0.05% TBST. Further blocking was done twice with 10% ADB (15 minutes each). Oocytes were then incubated overnight at 4°C with primary antibody: Mouse anti-TAp63α (Cat no. CM163B, 1:200). After incubation, oocytes were washed three times with 0.05% TBST, followed by two additional ADB blocking steps of 15 minutes each. The secondary antibody treatment involved incubation with Goat anti-mouse 555 (Cat no. A32727, Invitrogen, 1:1000) at 37°C for 1 hour in the dark. Following this, oocytes were washed thrice with 0.05% TBST. Oocytes were then mounted with prolong diamond (CAT No.- P36970, Invitrogen) and imaged.

Oocyte isolation was also done from late embryonic goat ovary (crown-rump length 30cm) and cultured in TCM199 (Cat. No. M2154, Sigma) with HIPK2 inhibitor (i_HIPK2-5 μM) and IKKβ inhibitor (i_IKKβ-5μM) along with control at 0^th^ hour, after 20 min DNA damage was introduced with doxorubicin (5 μM), following 6 hours of treatment oocytes were collected for RNA isolation using RNA isolation kit. (cat no.-12204-01, Thermo).

### 6.3 Fertility Studies

To assess fertility, adult female mice, including both treated and vehicle control groups, were individually housed with untreated males. If a female did not conceive within one month, the male was replaced. Mating success was monitored by checking for vaginal plugs, and if mating was unsuccessful, the female was paired with a new male until successful copulation occurred.

### 6.4 Endocrine Profiling

Serum samples were analysed for LH (Cat no.-KLM0573) and AMH (Cat no.-KLM1096) levels following the procedures outlined in the manufacturer’s protocol.

### 6.5 Oocyte Collection and Maturation

Ovarian puncture was performed using a 24-gauge needle in M2 medium (Cat. No. M7167, Sigma) to retrieve oocytes. The oocytes were carefully aspirated under a microscope, thoroughly washed, and subsequently cultured in M2 medium within a CO₂ incubator maintained at 37°C with 90–95% humidity. After 16 hours of incubation, oocyte maturation was assessed based on polar body formation and DNA staining.

### 6.6 Embryo Collection and Culture

Superovulation was induced in the animals using PMSG and HCG. The oviducts of plugged females were dissected and punctured with a 24-gauge needle in M2 medium to retrieve the embryos. The collected embryos were washed in M2 and EmbryoMax HTF medium (Cat. No. MR-070-D, Sigma) before being cultured in EmbryoMax HTF medium within a four-well IVF dish (Cat. No. 144444, Thermo). The culture was maintained in a CO₂ incubator at 37°C with 90–95% humidity. After four days, the embryos were evaluated for maturation.

### 6.7 RNA Isolation and mRNA Expression

Following euthanasia of the mice, the ovaries were preserved in 1 ml of Trizol (Cat No. 9108, Takara). The tissue was finely minced, vortexed, sonicated, and centrifuged at 10,000 rpm for 2 minutes at 4°C. The resulting supernatant was collected, and RNA extraction was performed using the phenol-chloroform method. RNA concentration was then measured, and cDNA synthesis was conducted according to the manufacturer’s protocol (Cat No. 6110a, Takara). For gene expression analysis, 100 ng of cDNA was used to assess the mRNA expression levels of *puma*, *noxa*, *bcl2l1*, *mcl1*, and *gapdh* (primer sequences provided in Table 1) in ovarian tissue. The analysis was performed using the TB Green SYBR mix (Cat No. RR820A, Takara), following the manufacturer’s instructions. Relative expression levels were quantified using the −ΔΔCt method as described by Livak, with statistical significance set at *p* < 0.05.

**Table 1:**
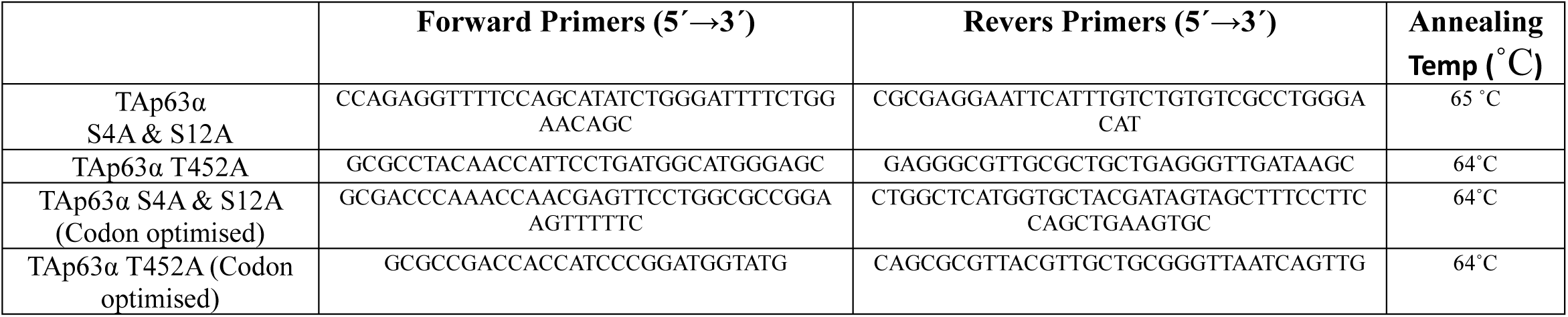
Primer list to generate TAp63α mutations.

### 6.8 Goat Ovary Culture

Ovary samples from Osmanabadi goats were collected from a slaughterhouse late fetal stage crown to rump 32cm and immediately immersed in warm saline (37°C) supplemented with penicillin–streptomycin antibiotics. The samples were transported to the laboratory within an hour using an embryo carrier. Upon arrival, the ovaries were trimmed and placed on a filter strip (Cat. No. 782810), which was floated on a nutrient medium containing TCM199 (Cat. No. M2154, Sigma), 10% goat follicular fluid, 1× penicillin–streptomycin (Cat. No. P4333, Sigma), and 3% fetal calf serum. The culture plates (Cat. No. 782891) were then incubated in a CO₂ incubator set at 37°C with 90–95% humidity. After two hours of culture, either 100 μM of i_HIPK2 (Cat. No. HY-U00439A, MedChem) or i_IKKβ (Cat. No. HY-13687, MedChem), along with their respective controls, were added. The culture was maintained for 48 hours, after which the ovaries were fixed in 10% formalin for further processing and sectioning. All experiments were performed in triplicate to ensure the reliability and consistency of findings.

### 7.1 Tissue Preparation and Immunohistochemistry

The ovaries were collected and fixed in 10% formalin for 2 to 4 days. Following fixation, they were subjected to a graded sucrose treatment using 10%, 20%, and 30% sucrose solutions for up to 3 days, depending on the ovary size. Cryosections of 5 μm thickness were prepared using the Cryostar NX50 and mounted onto positively charged glass slides. These slides were subsequently deparaffinized to prepare them for staining. Antigen retrieval was performed by immersing the slides in an antigen retrieval buffer (10 mM sodium citrate and 0.05% Tween 20) at 95°C for 60 minutes. The sections were then cooled to room temperature under running water, washed twice with deionized water (dH2O) for 5 minutes each, and rinsed in 1X TBST containing 0.1% Tween 20 for 1 minute. Blocking was carried out using M.O.M (BMK-2202, vector labs) if using mouse monoclonal primary antibody for mouse tissue for 1hr at RT followed by blocking with ADB twice for 15 mins each. The sections were incubated overnight with primary antibodies: [Mouse anti-TAp63α (Cat no. CM163B, 1:500), Rabbit anti-MVH (Cat no. ab13840, Abcam 1:500), Rabbit anti-Cleaved Caspase 3 (Cat no. PA5-114687, Thermo, 1:200), Mouse anti γH2A.X (05-636, Millipore,1:200)]. After incubation, the slides were washed twice with 1X TBST-0.1% Tween 20 for 10 minutes each, followed by two additional ADB blocking steps of 15 minutes each. The secondary antibody treatment involved incubation with Goat anti-mouse 555 (Cat no. A32727, Invitrogen, 1:2000) and Goat anti-rabbit 488 (Cat no. A32731, Invitrogen, 1:2000) at 37°C for 1 hour in the dark. Following this, the slides were washed three times with 1X TBST-0.1% Tween 20 for 10 minutes each and stained with 0.01% DAPI (Cat no. D9542, Sigma) for 2 minutes. After a brief rinse in dH2O, the slides were air-dried. For DAB staining, after the ADB blocking step, the slides were incubated with Goat anti-mouse biotin (Cat no. A16076, Invitrogen, 1:1000), Goat anti-rabbit biotin (B-2770, Invitrogen,1:1000) at 37°C for 1 hour in the dark. The slides were then washed three times with 1X TBST-0.1% Tween 20 for 5 minutes each, followed by processing according to the manufacturer’s instructions using the VECTASTAIN ABC-HRP kit (Cat no. PK-6100, vector labs) and the DAB substrate kit (Cat no. SK-4100, vector labs). Finally, the slides were dried, mounted with DPX, and imaged using a microscope as we previously published (Mohanty et al., 2024).

### 7.2 Imaging

Immunofluorescence-stained ovaries were imaged using a Zeiss Axio scope VII microscope equipped with 10×, 20×, or 63× Plan Apochromat objectives, an EXFO X-Cite metal halide light source, and a Hamamatsu ORCA-ER CCD camera. Images were processed with Zen software. Immunohistochemistry-stained ovary sections were imaged with a Nikon Eclipse Ti2 Inverted Microscope and processed with NIS-Elements software.

### 7.3 Quantification

Every fifth 5μm section from each ovary was analysed to assess follicular development stages, including primordial, primary, secondary, preantral, and antral follicles. Follicles were classified based on the number of cell layers and the presence of an antrum. To minimize bias, a standardized counting method was applied consistently across all groups. Given the impracticality of counting all follicles from every fifth section in goats, a single middle section from each of the three experimental replicate datasets was examined to quantify follicles. All quantitative analyses were performed by two observers, one of whom was blinded to the experimental groups.

### 8.1 Western blotting

Cell pellets or ovaries were resuspended in a lysis buffer consisting of 25 mM Tris-HCl (pH 8.0), 250 mM NaCl, 1% Triton X-100, 0.1% SDS, 2 mM MgCl₂, PMSF, protease inhibitor, phosphatase inhibitor, and sodium-β-glycerophosphate. The samples were then subjected to sonication and centrifuged at 15,000 rpm for 10 minutes at 4°C. The resulting supernatants were collected and subsequently boiled at 100°C for 5 minutes in 1X Laemmli buffer before being loaded onto an SDS-PAGE gel for protein analysis. Following electrophoresis, proteins were transferred onto a nitrocellulose membrane (Bio-Rad) and blocked with 5% skimmed milk for 1 hour at room temperature. The membrane was then incubated overnight at 4°C with primary antibodies, including Mouse anti-TAp63α (Cat. No. CM-163B, 1:1000) and Mouse anti-β-Actin (Cat. No. SC-47778, 1:1000). After incubation, the membrane was washed three times with 1X TBST containing 0.3% Tween 20, each wash lasting 5 minutes. It was then incubated with HRP-conjugated secondary antibodies (Goat anti-Mouse IgG HRP, Cat. No. 31430) for 2 hours at room temperature. Following secondary antibody incubation, the membrane was washed three more times with 1X TBST-0.3% Tween 20 for 5 minutes each, followed by a final wash with 1X TBS. The blot was subsequently developed using a chemiluminescence detection method.

### 8.2 Western blot quantification

Western blot band intensities were quantified using ImageJ, ensuring high-resolution images for precise analysis. The rectangular selection tool was used to define the region of interest (ROI) by drawing a box around each band, ensuring it was slightly larger than the band while excluding background noise. This ROI was saved, and the same box size was applied to all bands for consistency. Each band was added to the ROI Manager, and its intensity was measured, generating a results table. Background subtraction was performed by measuring the intensity of a blank area near the bands and subtracting this value from all band intensities. Target band intensities were normalized to the corresponding loading control, β-actin, by dividing the intensity of each target band by that of the control band. For phosphorylated protein quantification, the intensity of the phosphorylated band was normalized to the total protein. The normalized data were analysed in GraphPad Prism to determine relative expression levels or fold changes. Consistent protein loading and image quality were maintained to ensure accurate quantification.

## 9. STRING analysis

Proteins identified from mass spectrometry analysis were used as input for the STRING database to perform protein-protein interaction analysis. The identified proteins were uploaded to the STRING database. A confidence score threshold was applied to filter the interactions, with higher thresholds ensuring more reliable evidence. The interaction network was visually analysed to identify key hub proteins and exported. Care was taken to validate the input data and configure STRING parameters to align with the research objectives.

## Methods contact

Further information and requests for resources and reagents should be directed to the lead contact, H.B.D. Prasada Rao

## ACKNOWLEDGMENTS

We thank the NIAB core Microscope and small animal facility. A.K. was supported by DBT SRF. A.M. was supported by CSIR SRF. S.A. was supported by UGC JRF. L.K.S was supported by DBT SRF. A.K., M.A. was supported by UGC SRF. P.B., A.K., S.A. was supported by UGC JRF. This work was supported by NIAB core grant C0031 awarded to H.B.D.P.R.

## AUTHOR CONTRIBUTIONS

A.K., and H.B.D.P.R. conceived the study and designed the experiments. A.K., P.K., A.M., S.A. and H.B.D.P.R. performed the experiments and analyzed the data. H.B.D.P.R. and A.K. wrote the manuscript with inputs and edits from all authors.

## Data availability

The data that support the findings of this study are available from the corresponding author upon request.

## Supplementary Figure Legends

**Model:**

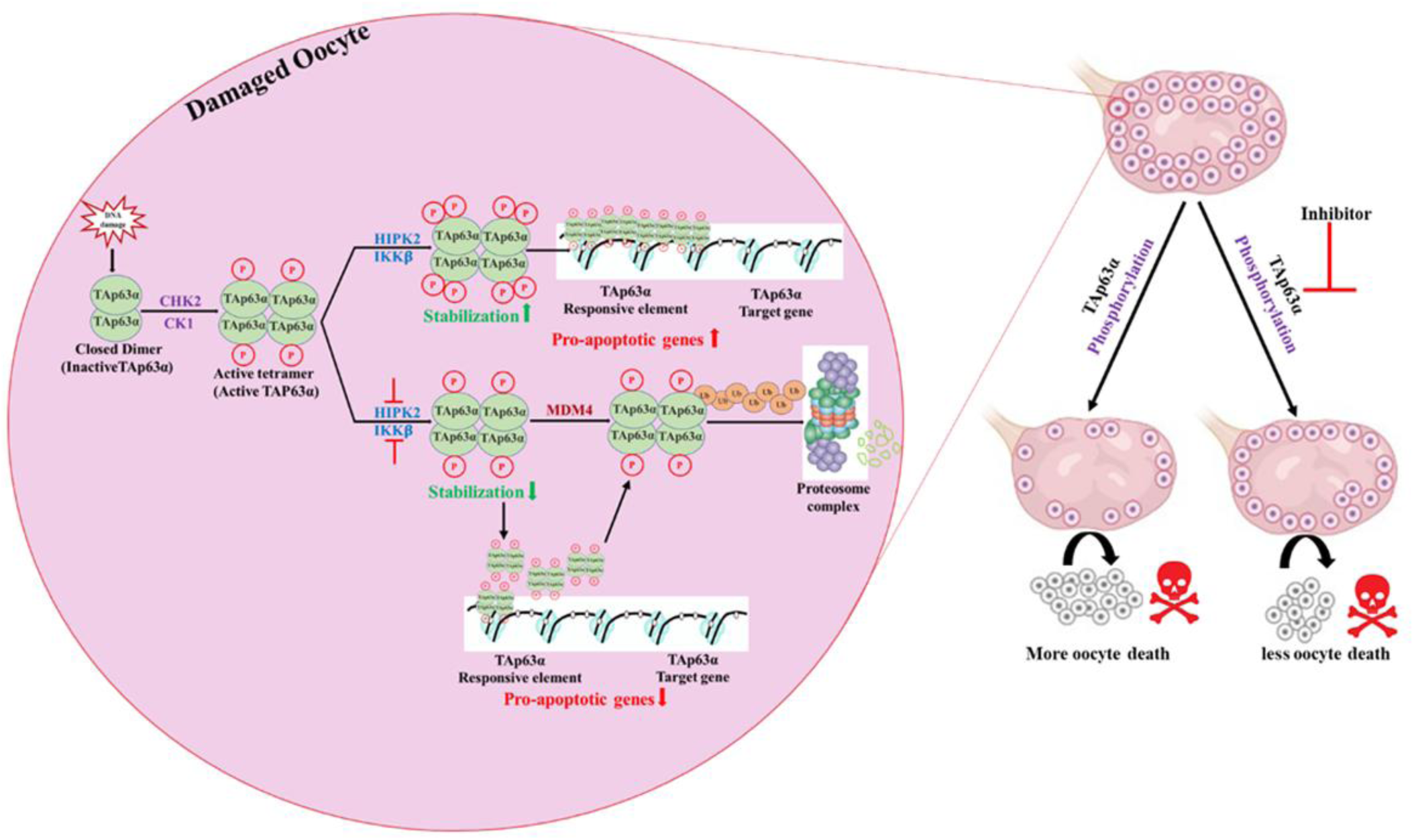

**Model:**

Under genotoxic stress, a coordinated kinase signaling network regulates the stability of TAp63α, a key guardian of the early follicles. CHK2 initiates phosphorylation, followed by CK1-mediated modifications that facilitate tetramerization. Concurrently, HIPK2 and IKKβ phosphorylate TAp63α at threonine 452, serine 4, and serine 12, preventing MDM4-mediated ubiquitination and subsequent proteasomal degradation. This stabilization mechanism ensures the integrity of oocytes, even in the presence of DNA-damaging agents such as cisplatin. Notably, the pharmacological inhibition of HIPK2 and IKKβ leads to significant protection of the follicles, highlighting their critical role in follicle pool maintenance. Collectively, multi-kinases orchestrate a phosphorylation-dependent regulatory pathway that safeguards oocytes under genotoxic stress.

**Figure S1.**
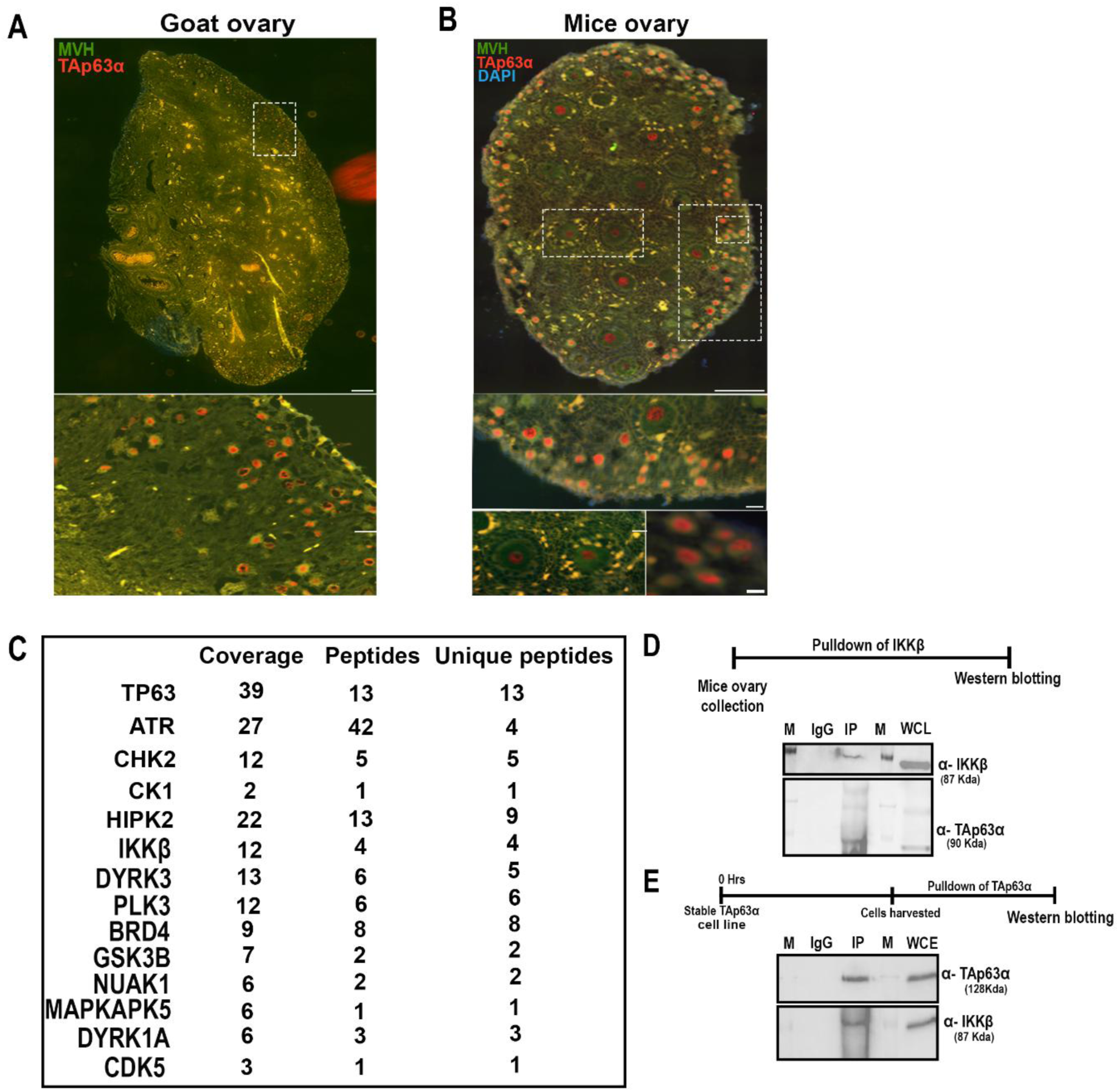
(A) Immunofluorescence image of goat ovary sections stained with MVH (green) and TAp63α (red), validating the specificity of the TAp63α antibody. A magnified view of the selected region is shown below. (B) Immunofluorescence image of mouse ovary sections stained with MVH (green) and TAp63α (red). A magnified view of the selected region is shown below. (C) Table summarizing proteins identified via mass spectrometry following TAp63α immunoprecipitation (IP). The table includes protein coverage, total peptides detected, and uniquely identified peptides. (D) Immunoprecipitation of IKKβ from mouse ovary, followed by detection using anti-IKKβ and anti-TAp63α antibodies. (E) Western blot analysis validating the pulldown of TAp63α from the stable TAp63α cell line. Detection was performed using anti-TAp63α antibodies for TAp63α and anti-IKKβ for IKKβ respectively.

**Figure S2.**
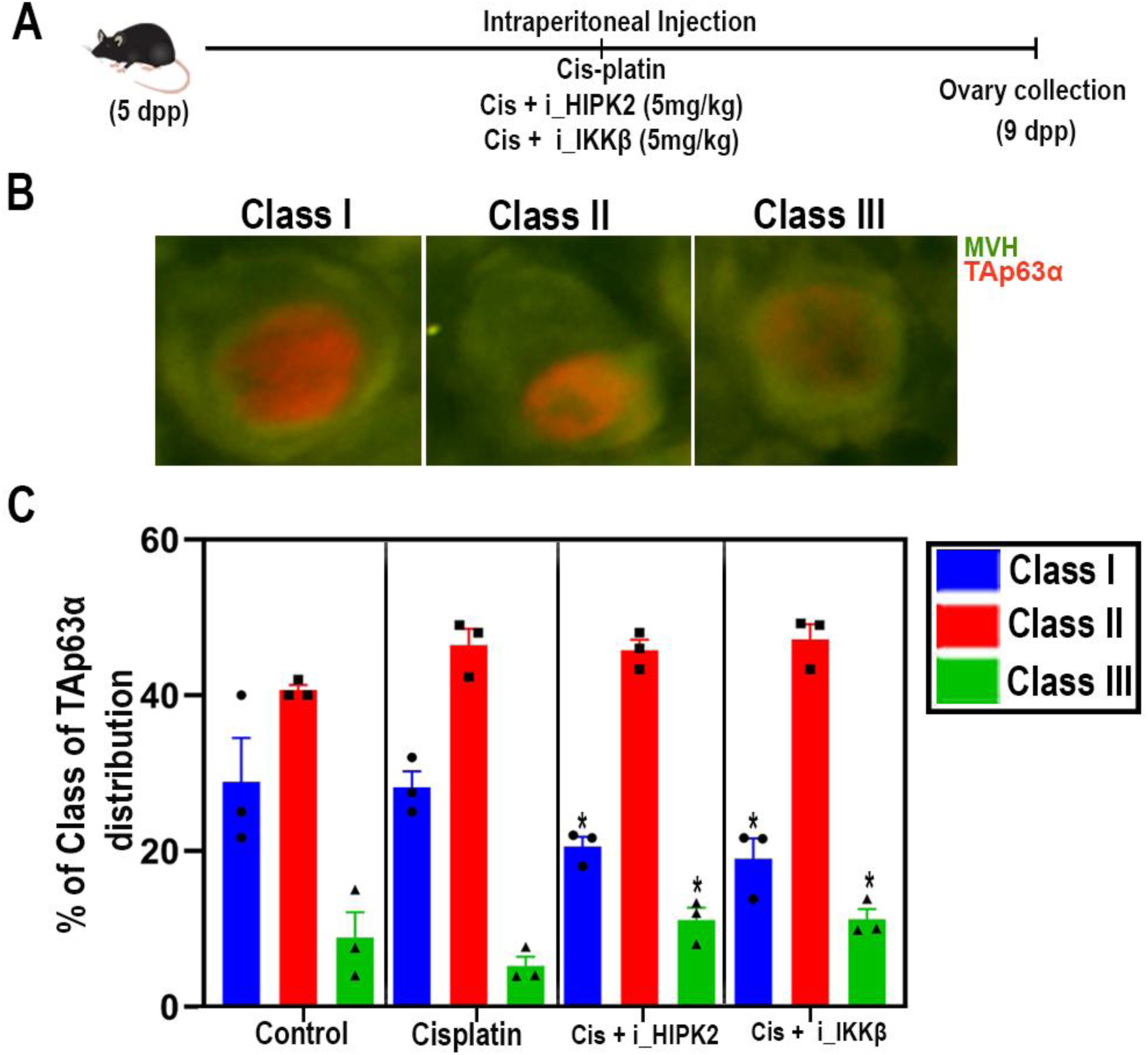
(A) Schematic representation of the experimental workflow. (B, C) Image shows the different class of TAp63α distribution and their quantification. Statistical significance for class III follicles: Cis + HIPK2 inhibitor (*P = 0.04*), Cis + IKKβ inhibitor (*P = 0.03* and for class I follicles, Cis + HIPK2 inhibitor (*P = 0.03*), Cis + IKKβ inhibitor (*P = 0.04*) in comparison to only cisplatin group. *unpaired* t*-test,* Data are presented as mean ± SD.

**Figure S3.**
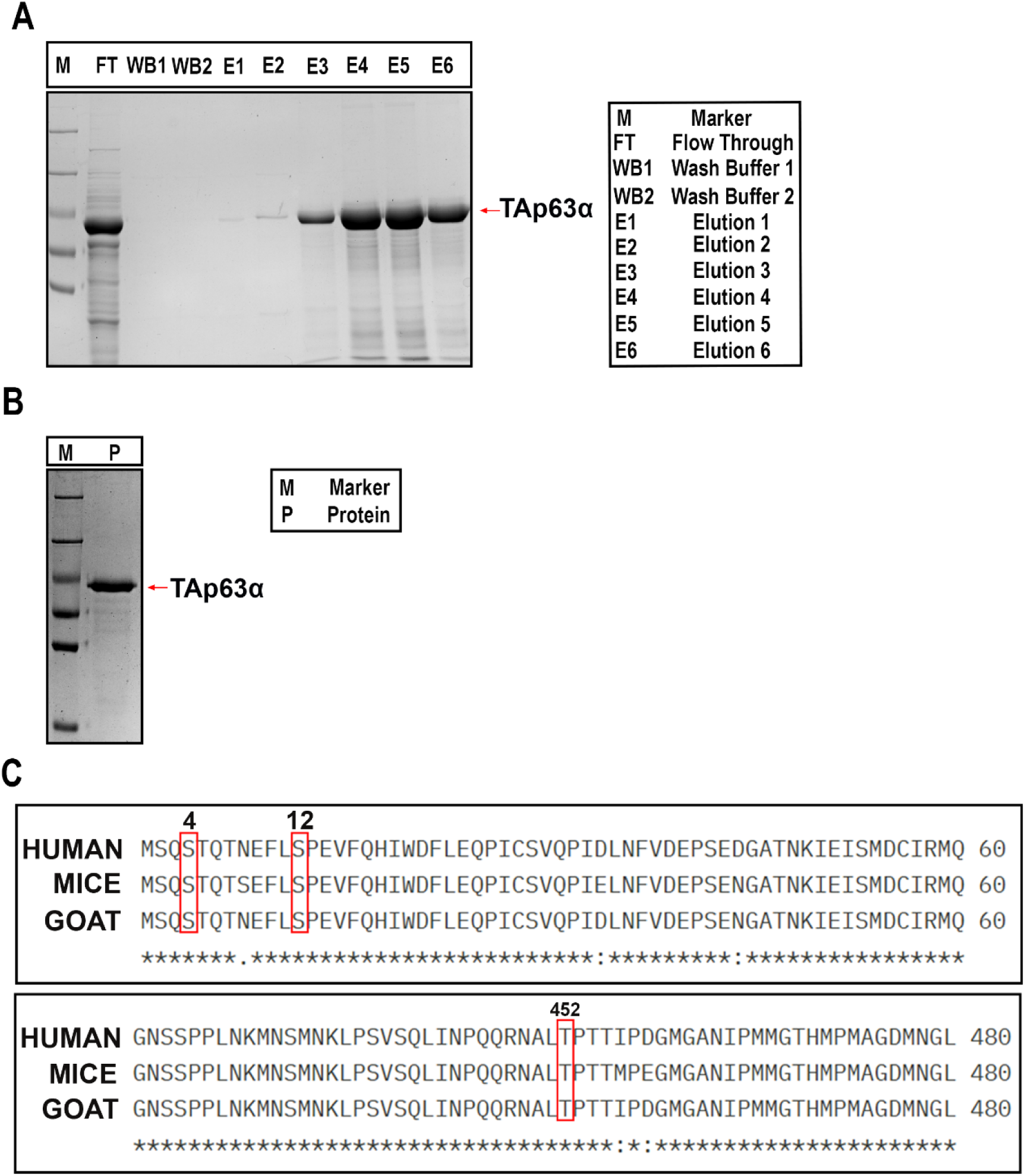
(A) Purification of recombinant TAp63α protein from *E. coli* after transforming plasmid containing TAp63α gene. Eluted with different concentrations of imidazole 50mM, 100mM, 200mM, 300mM, 400mM, 500mM). (B) Purified recombinant TAp63α protein after dialysis in Tris (20mM) and NaCl (100mM). (C) Multiple sequence alignment of TP63 from human, mouse and goat, shows conserved phosphorylation sites targeted by HIPK2 (T452) and IKKβ (S4 and S12).

**Figure S4.**
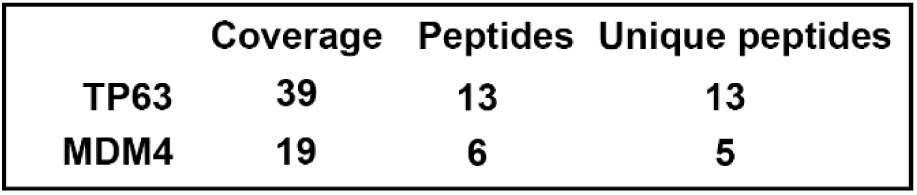
Table summarizing proteins identified via mass spectrometry including protein coverage, total peptides detected, and uniquely identified peptides.

**Figure S5.**
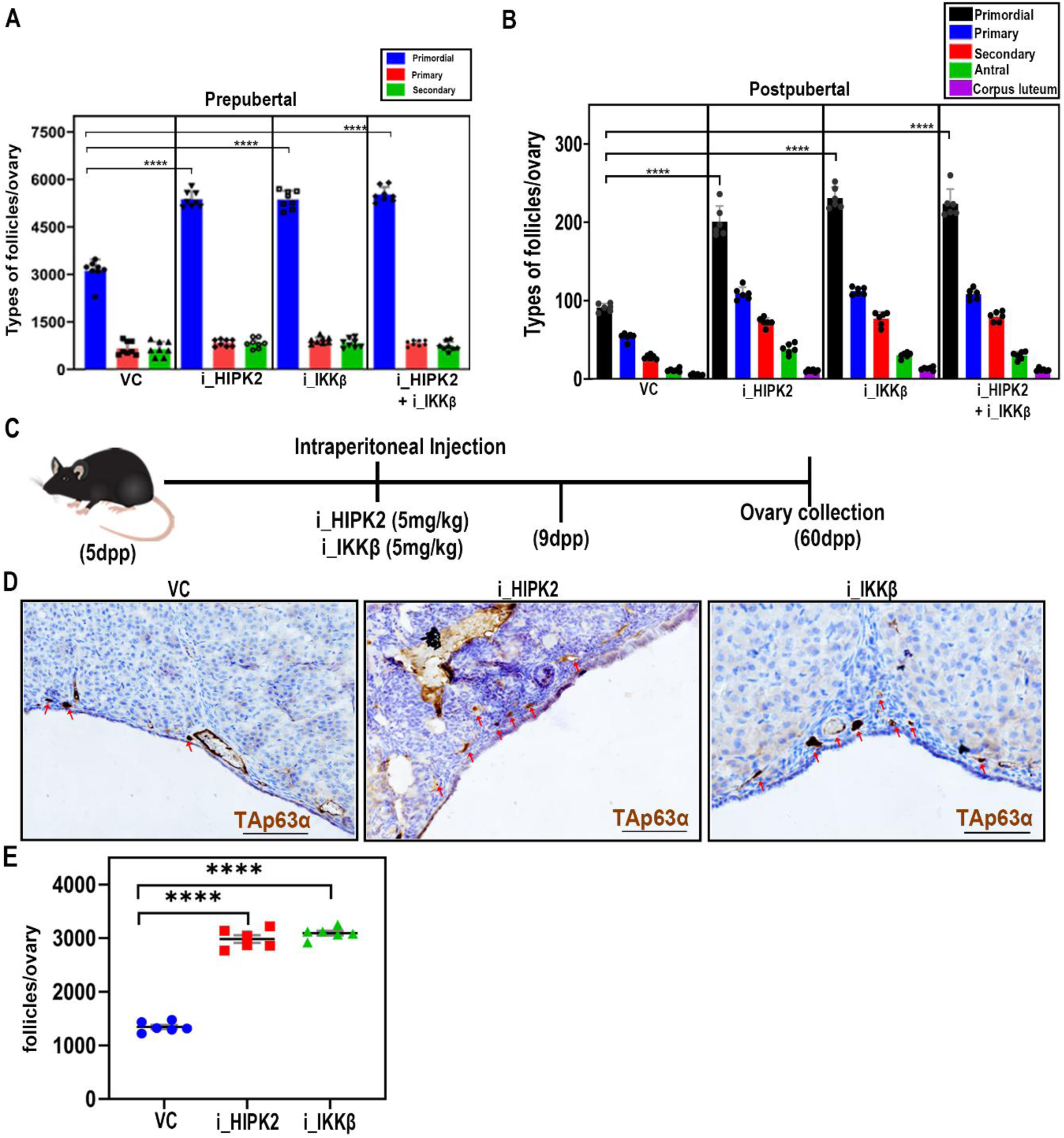
(A) Quantification of primordial, primary, and secondary follicles at 9 days post-partum (dpp) in vehicle control (VC), HIPK2 inhibitor (i_HIPK2), IKKβ inhibitor (i_IKKβ), and the combination of both inhibitors (i_HIPK2 + i_IKKβ). Statistical analysis of primordial follicle counts shows significant differences with P-values as follows: i_HIPK2 \*\*\*\**P ≤ 0.0001*, i_IKKβ \*\*\*\**P ≤ 0.0001*, and i_HIPK2 + i_IKKβ \*\*\*\**P ≤ 0.0001* (B) Schematic representation of the experimental design for long-term ovarian assessment. (C) Representative images of ovarian sections from 60dpp C57BL/6 mice treated with Vehicle control, HIPK2 inhibitor and IKKβ inhibitor. Ovarian sections were immunostained for TAp63α (brown) with hematoxylin counterstaining. (D) Quantification of the total follicles from panel. Statistical analysis indicates significant preservation of follicles in HIPK2 inhibitor (*P ≤ 0.0001*), IKKβ inhibitor (*P ≤ 0.0001*) compared to the Vehicle control group. (E) Quantification of primordial, primary, and secondary follicles at 5 months in VC, i_HIPK2, i_IKKβ, and i_HIPK2 + i_IKKβ treatment groups. Statistical analysis of primordial follicle counts shows significant differences with P-values as follows: i_HIPK2 \*\*\*\**P ≤ 0.0001*, i_IKKβ \*\*\*\**P ≤ 0.0001*, and i_HIPK2 + i_IKKβ \*\*\*\**P ≤ 0.0001*. Data are presented as mean ± SD, and error bars indicate standard deviation Statistical significance was determined using an *unpaired t-test*. Scale bars for 60 days mice ovary section: 50μm.

**Figure S6.**
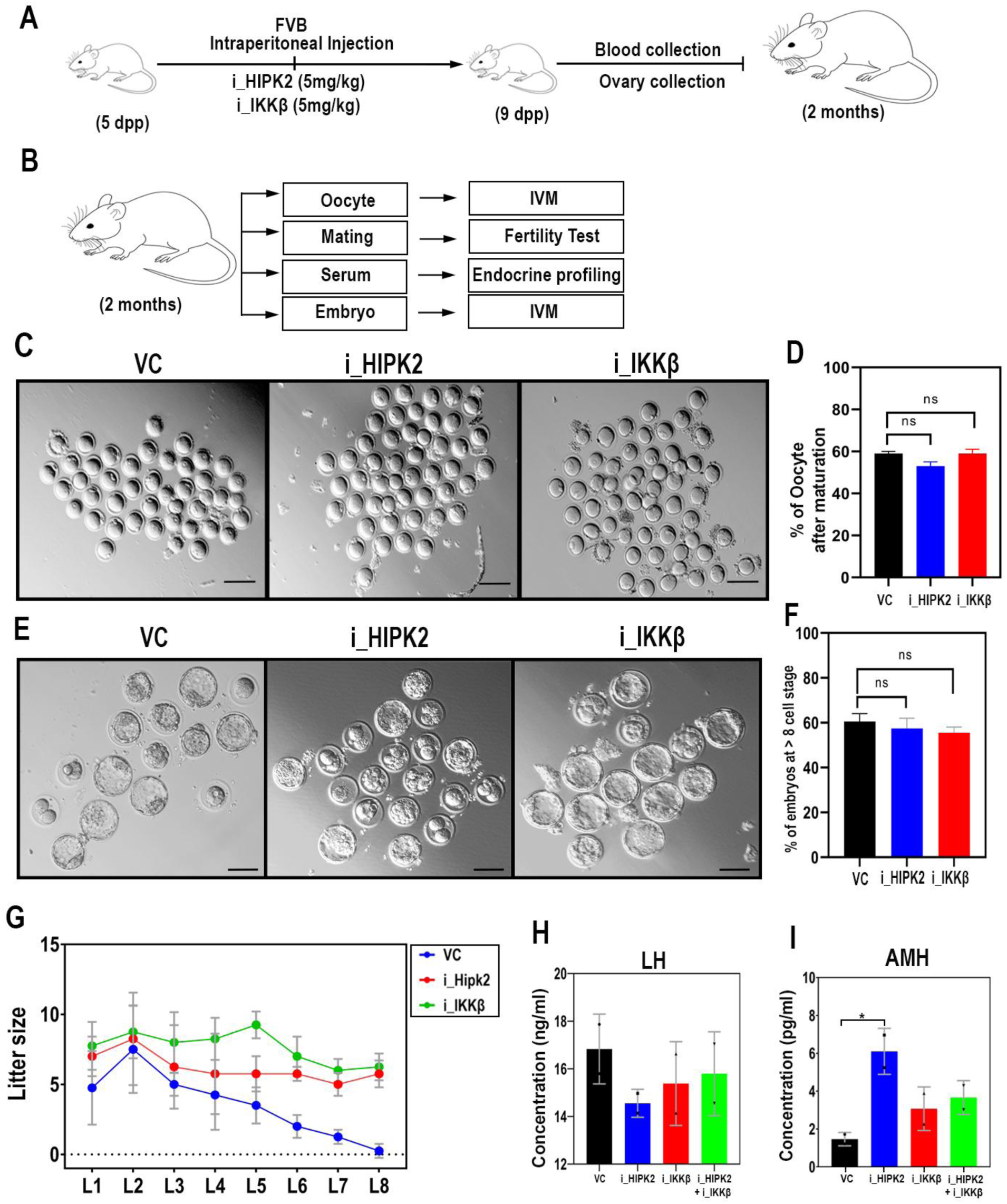
(A) Schematic representation of the experimental protocol for intraperitoneal injection of Vehicle Control (VC), HIPK2 inhibitor, and IKKβ inhibitor, followed by (B) oocyte collection, fertility testing, and endocrine profiling. (C) Bright-field images showing oocyte maturation across different treatment groups. (D) Quantification of in vitro oocyte maturation in females treated with VC, HIPK2 inhibitor, IKKβ inhibitor. (E) Bright field images of embryo development (F) Quantification of in-vitro embryo development from VC, HIPK2 inhibitor, IKKβ inhibitor. (G) Fertility assessment of females treated with VC, HIPK2 inhibitor, and IKKβ inhibitor, and a combination of HIPK2 and IKKβ inhibitors. (H, I) Quantification of serum LH and AMH levels across treatment groups. Statistical significance was determined using an *unpaired t-test*. Data are presented as mean ± SD. Non-significant p-values are not displayed. Scale bars: 200 μm.

**Figure S7.**
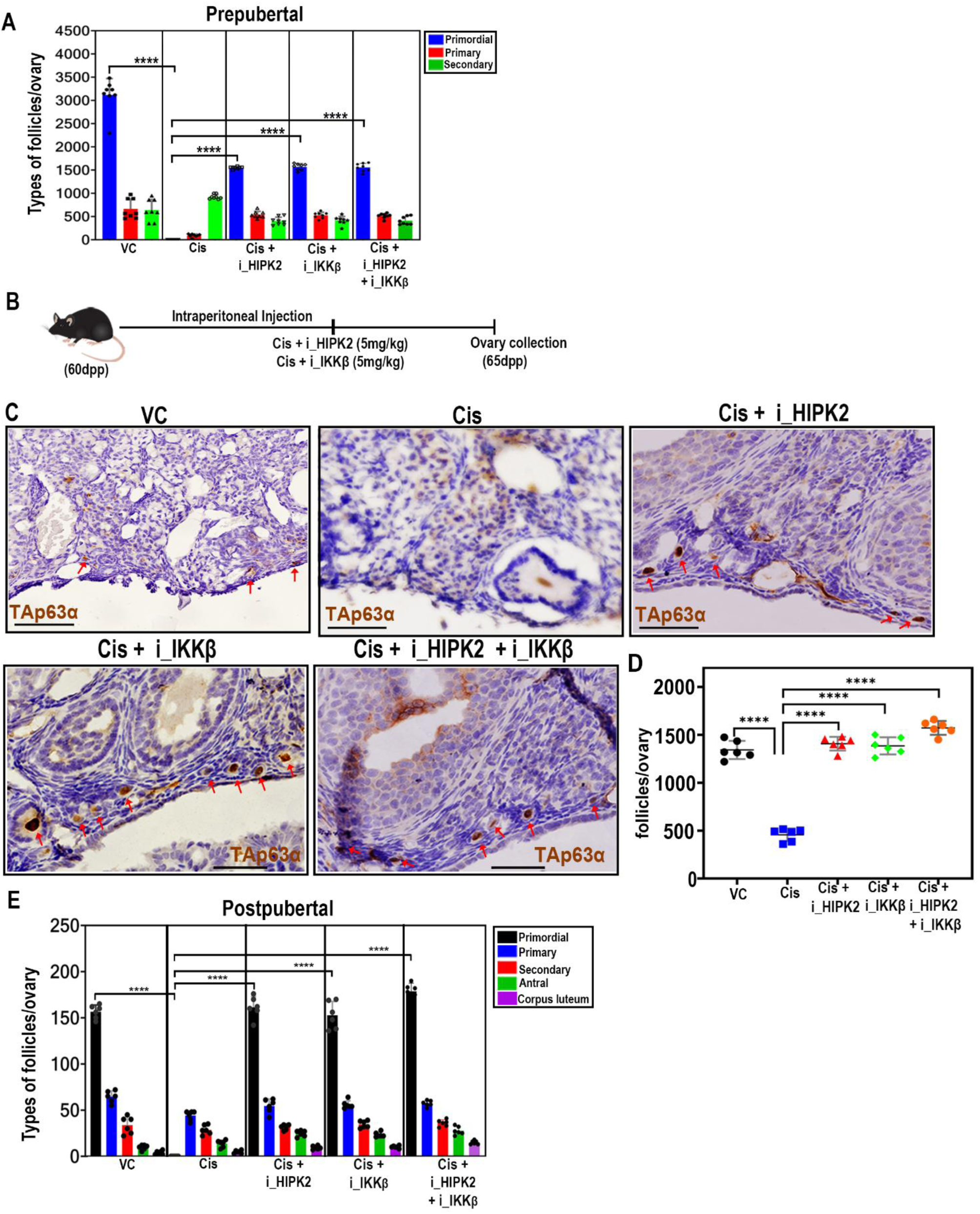
(A) Quantification of primordial, primary, and secondary follicles at 9 days post-partum (dpp) in ovaries from mice treated with Vehicle control, Cisplatin, Cisplatin + HIPK2 inhibitor, Cisplatin + IKKβ inhibitor, and Cisplatin + HIPK2 & IKKβ inhibitors. Statistical analysis difference in primordial follicle counts compared to the cisplatin group with P-values as follows: Vehicle control \*\*\*\**P ≤ 0.0001*, Cis + i_HIPK2 \*\*\*\**P ≤ 0.0001*, Cis + i_IKKβ \*\*\*\**P ≤ 0.0001*, and Cis + i_HIPK2 + i_IKKβ \*\*\*\**P ≤ 0.0001* (B) Schematic representation of the experimental design for long-term ovarian assessment under genotoxic stress. (C) Representative images of ovarian sections from 65dpp C57BL/6 mice treated with Vehicle control, Cisplatin, Cisplatin + HIPK2 inhibitor, Cisplatin + IKKβ inhibitor, Cisplatin +HIPK2+IKKβ inhibitor. Ovarian sections were immunostained for TAp63α (brown) with hematoxylin counterstaining. (D) Quantification of the follicles from panel. Statistical analysis indicates significant difference in Vehicle control (*P ≤ 0.0001*), Cisplatin + HIPK2 inhibitor (*P ≤ 0.0001*), Cisplatin + IKKβ inhibitor (*P ≤ 0.0013*), and Cisplatin + combined inhibitors (*P ≤ 0.0001*) compared to the Cisplatin-only. (E) Quantification of primordial, primary, and secondary follicles, Statistical analysis of primordial follicle counts shows significant differences with P-values as follows: Vehicle control \*\*\*\**P ≤ 0.0001*, Cis + i_HIPK2 \*\*\*\**P ≤ 0.0001*, Cis + i_IKKβ \*\*\*\**P ≤ 0.0001*, and Cis + i_HIPK2 + i_IKKβ \*\*\*\**P ≤ 0.0001*. Statistical analysis was performed using an *unpaired t-test*. Data are presented as mean ± SD. Non-significant P-values are not displayed in the figure. Scale bars for 9dpp mice ovary section: 100μm and for 60 days mice ovary section: 50μm.

**Figure S8.**
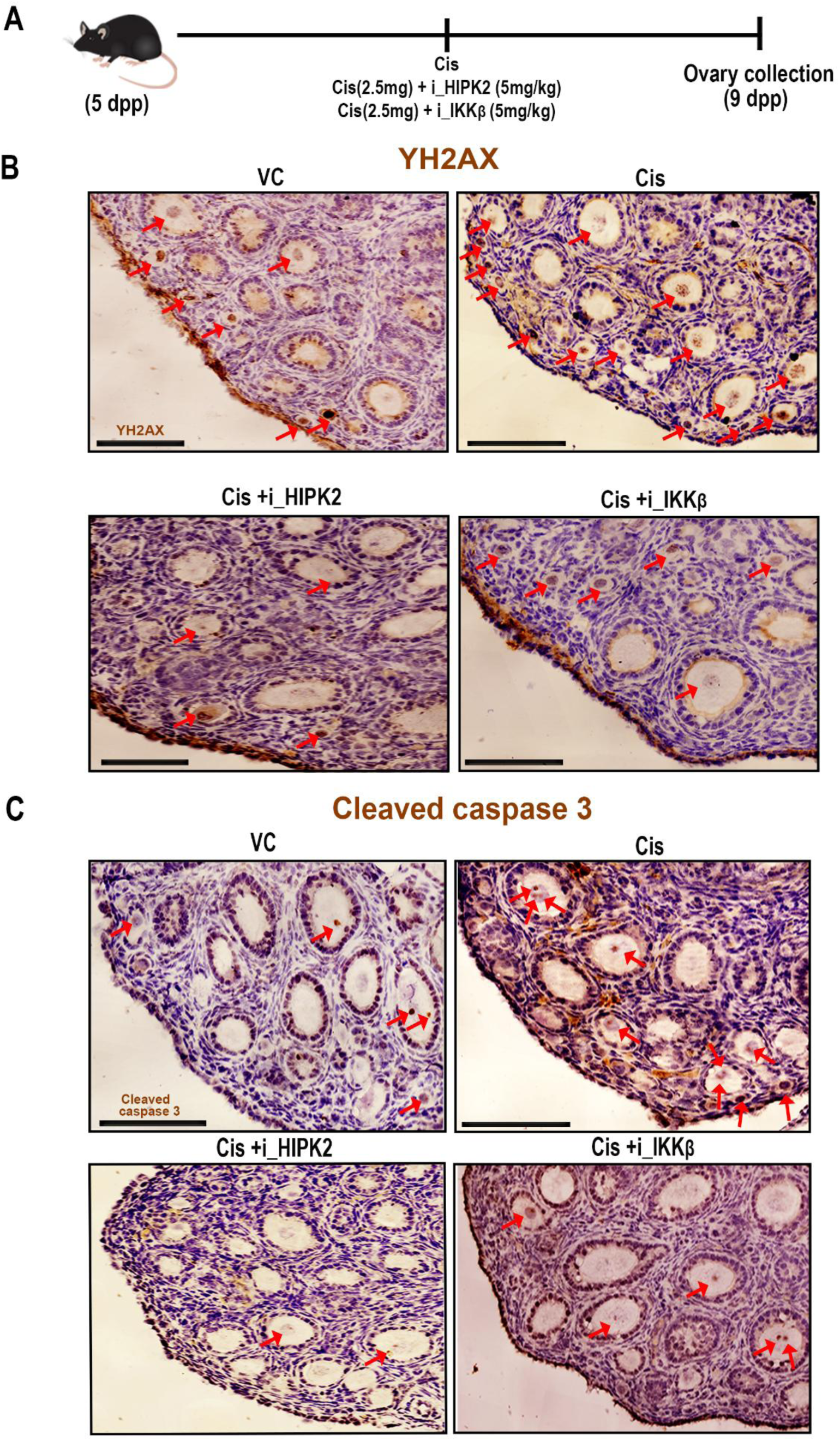
(A) Schematic representation of the experimental workflow. Representative images of ovarian sections from 9-day-postpartum (dpp) C57BL/6 mice treated with Vehicle control, Cisplatin, Cisplatin + HIPK2 inhibitor, Cisplatin + IKKβ inhibitor. Ovaries were immunostained for γH2AX (brown) (B) and for cleaved caspase3 (C) (brown) with hematoxylin counterstaining.

**Figure S9.**
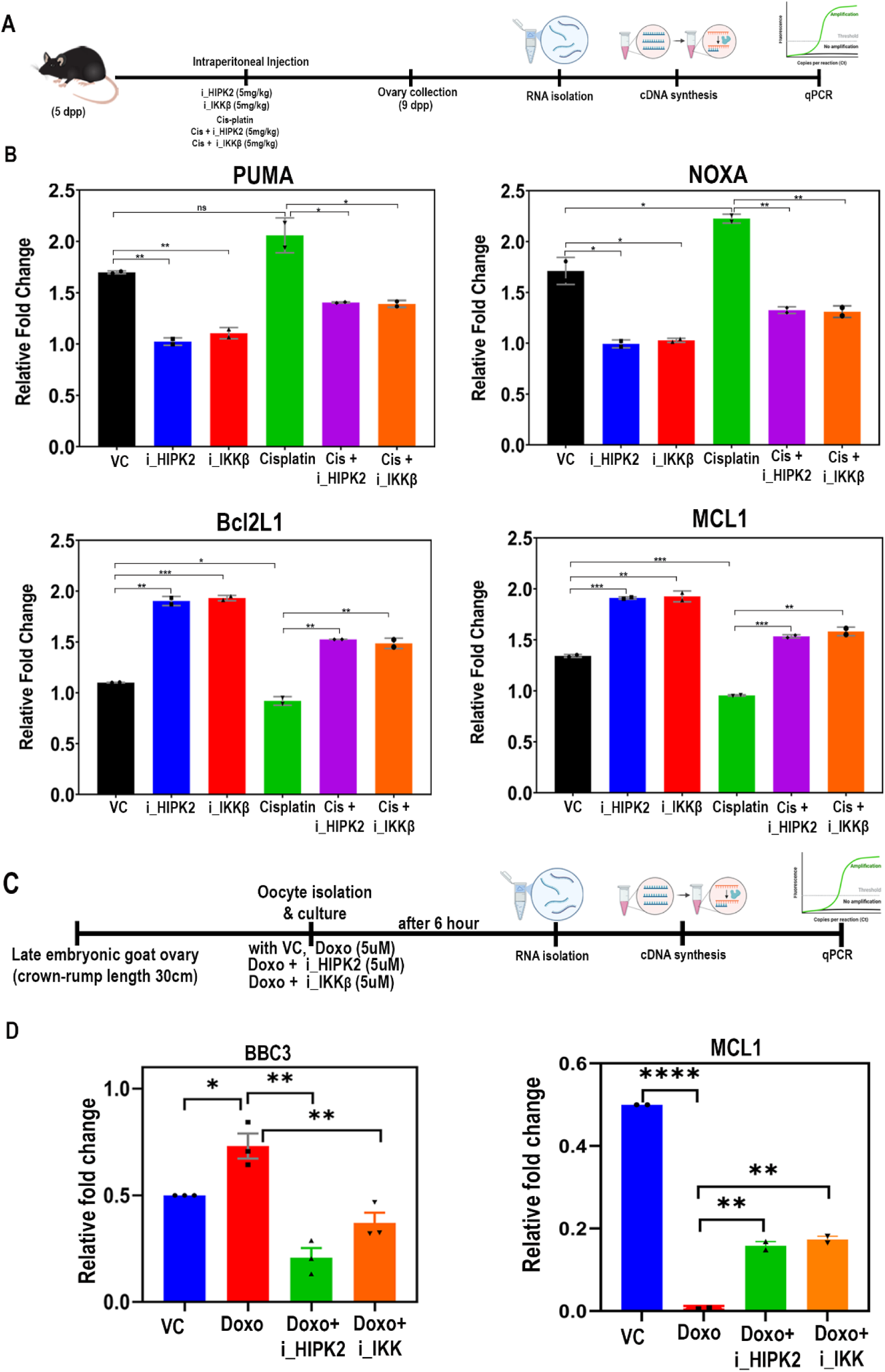
(A) Schematic representation of the experimental design for intraperitoneal administration of vehicle control (VC), HIPK2 inhibitor, and IKKβ inhibitor under normal and DNA damage conditions. Following treatment, ovarian tissues were collected for mRNA isolation and subsequent RT-PCR analysis. (B) Quantification of the relative fold change in expression of pro-apoptotic genes (*Puma* and *Noxa*) and anti-apoptotic genes (*Bcl2l1* and *Mcl1*) across different treatment conditions, including VC, HIPK2 inhibitor, IKKβ inhibitor, cisplatin, cisplatin + HIPK2 inhibitor, and cisplatin + IKKβ inhibitor. Statistical analysis of gene expression changes with unpaired *t*-tests. The *P* values are as follows: Puma: VC vs. HIPK2 inhibitor (*P ≤ 0.0016*), VC vs. IKKβ inhibitor (*P ≤ 0.0045*), VC vs. cisplatin (ns), cisplatin vs. cisplatin + HIPK2 inhibitor (*P ≤ 0.0235*), cisplatin vs. cisplatin + IKKβ inhibitor (*P ≤ 0.0236*). Noxa: VC vs. HIPK2 inhibitor (*P ≤ 0.0181*), VC vs. IKKβ inhibitor (*P ≤ 0.0188*), VC vs. cisplatin (*P ≤ 0.035*), cisplatin vs. cisplatin + HIPK2 inhibitor (*P ≤ 0.0018*), cisplatin vs. cisplatin + IKKβ inhibitor (*P ≤ 0.0031*). Bcl2L1: VC vs. HIPK2 inhibitor (*P ≤ 0.0016*), VC vs. IKKβ inhibitor (*P ≤ 0.0005*), VC vs. cisplatin (*P ≤ 0.0264*), cisplatin vs. cisplatin + HIPK2 inhibitor (*P ≤ 0.0024*), cisplatin vs. cisplatin + IKKβ inhibitor (*P ≤ 0.0067*). Mcl1: VC vs. HIPK2 inhibitor (*P ≤ 0.0005*), VC vs. IKKβ inhibitor (P ≤ 0.0044), VC vs. cisplatin (*P ≤ 0.0008*), cisplatin vs. cisplatin + HIPK2 inhibitor (*P ≤ 0.0004*), cisplatin vs. cisplatin + IKKβ inhibitor (*P ≤ 0.0023*). (C) Schematic representation of the experimental workflow. (D) Quantification of the relative fold change in expression of pro-apoptotic genes (*Bbc3*) and anti-apoptotic genes (*Mcl1*) across different treatment conditions, including VC, Doxo, Doxo + HIPK2 inhibitor, and Doxo + IKKβ inhibitor. The *P* values are as follows: BBC3: VC vs. Doxo (*P ≤ 0.017*), Doxo vs. Doxo + HIPK2 inhibitor (*P ≤ 0.0091*), Doxo vs. Doxo + IKKβ inhibitor (*P ≤ 0.0021*. MCL1: VC vs. Doxo (*P ≤ 0.0001*), Doxo vs. Doxo + HIPK2 inhibitor (*P ≤ 0.0044*), Doxo vs. Doxo + IKKβ inhibitor (*P ≤ 0.0024*). Statistical analysis was performed using an *unpaired t-test*. Data are presented as mean ± SD.

**Figure S10.**
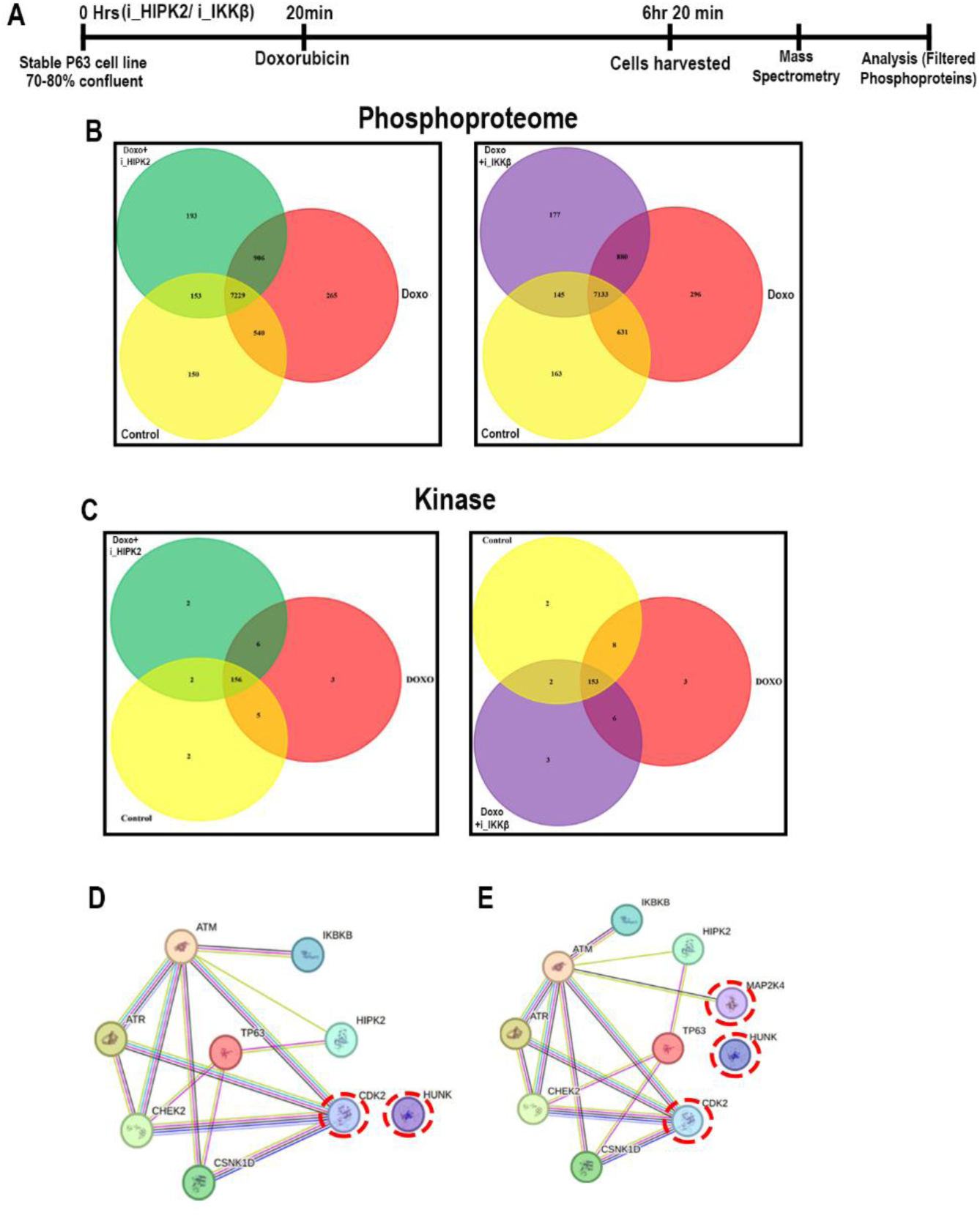
(A) Schematic representation of the experimental approach. Proteomic profiling of stable TAp63α-expressing H1299 cells treated with Doxo, Doxo + i_Hipk2, Doxo + i_IKKβ along with control. (B) Venn diagram of total phospho-proteome of control, Doxo and Doxo + i_Hipk2 & control, Doxo and Doxo + i_IKKβ (C) Venn diagram of total kinases identified from mass spec analysis in control, Doxo and Doxo + i_Hipk2 & control, Doxo and Doxo + i_IKKβ. STRING analysis illustrating the predicted interactions of TAp63α with kinases specifically present in Doxo + i_Hipk2 (D) and Doxo + i_IKKβ (E) from mass spec.

**Figure S11.**
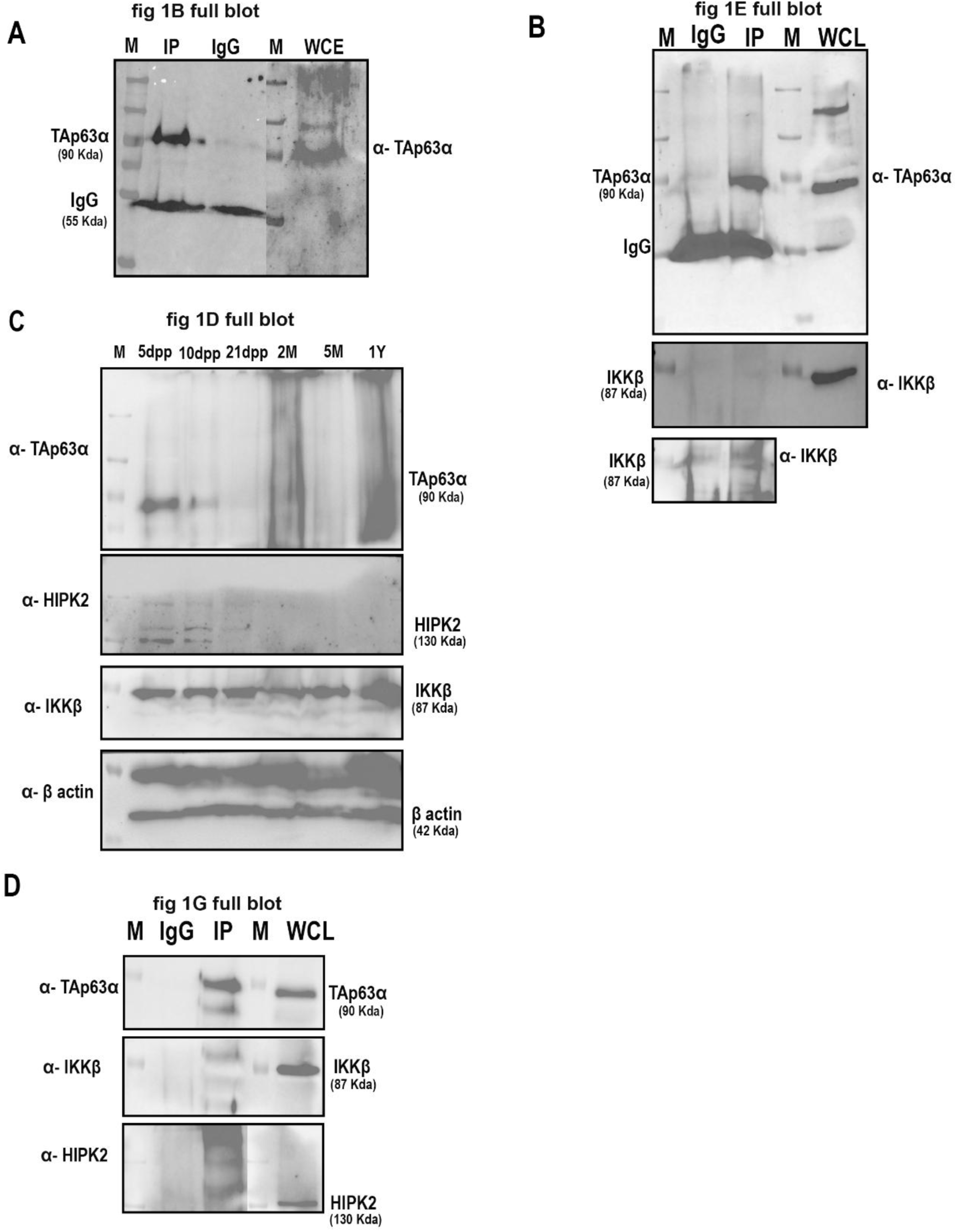
Whole western blots.

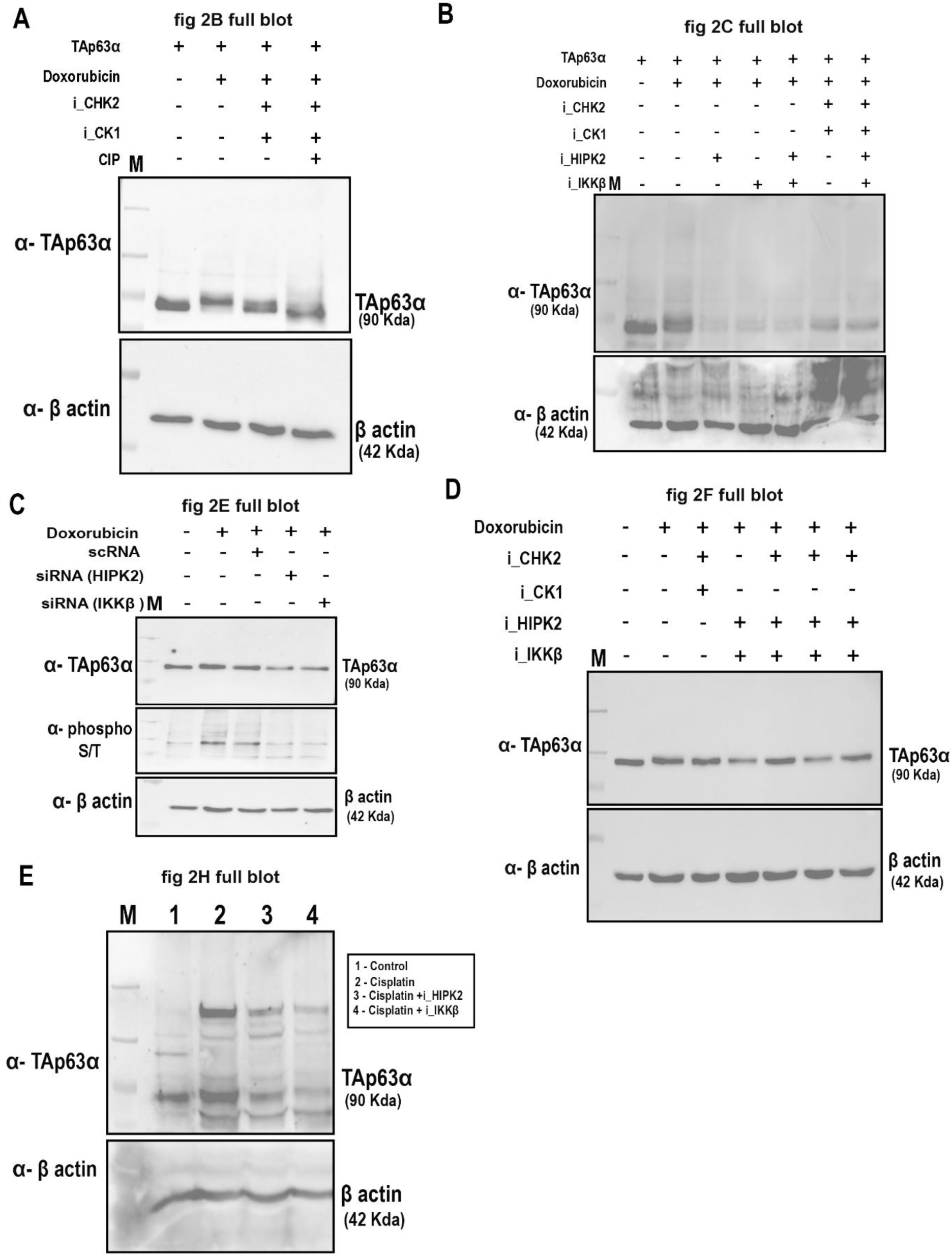

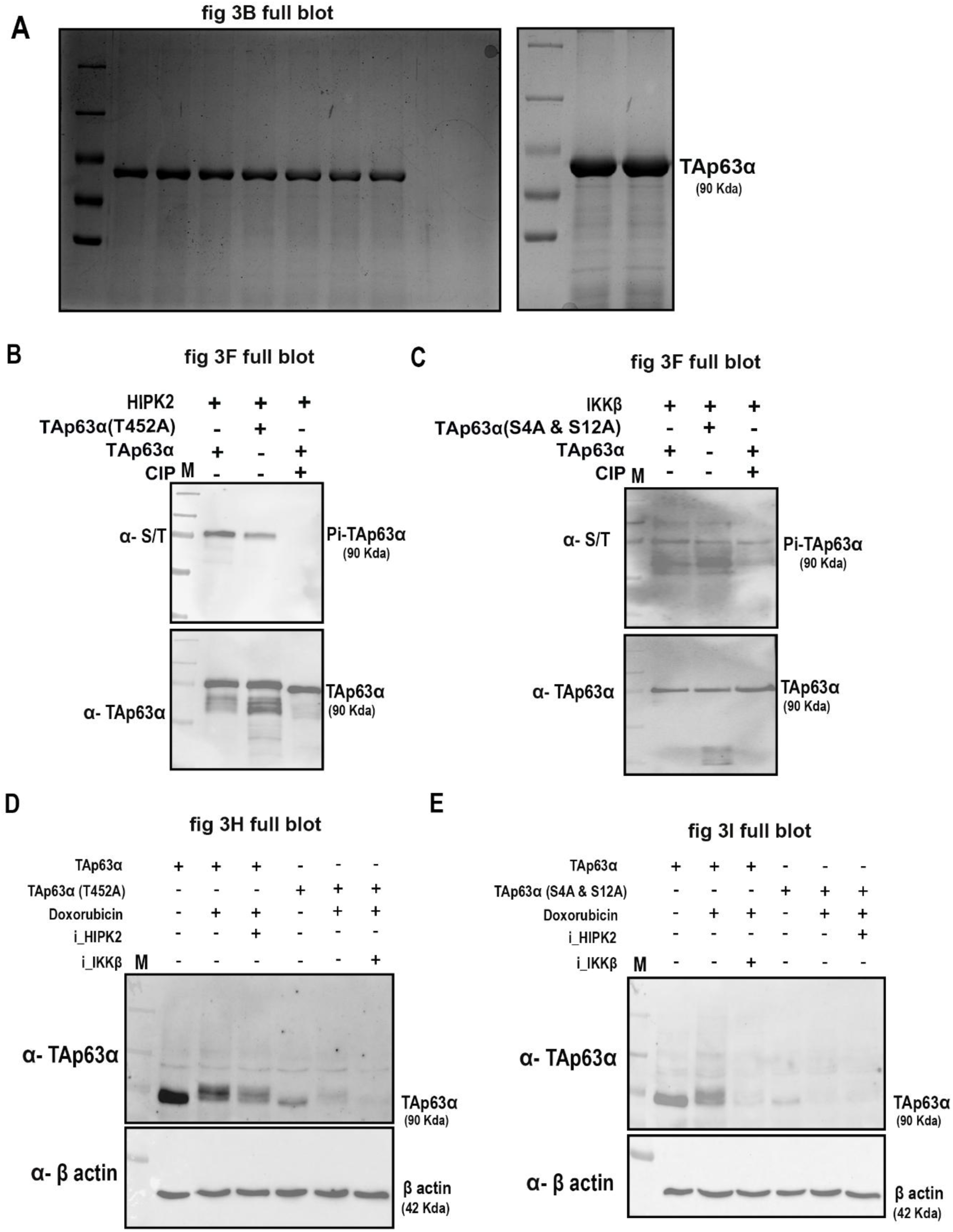

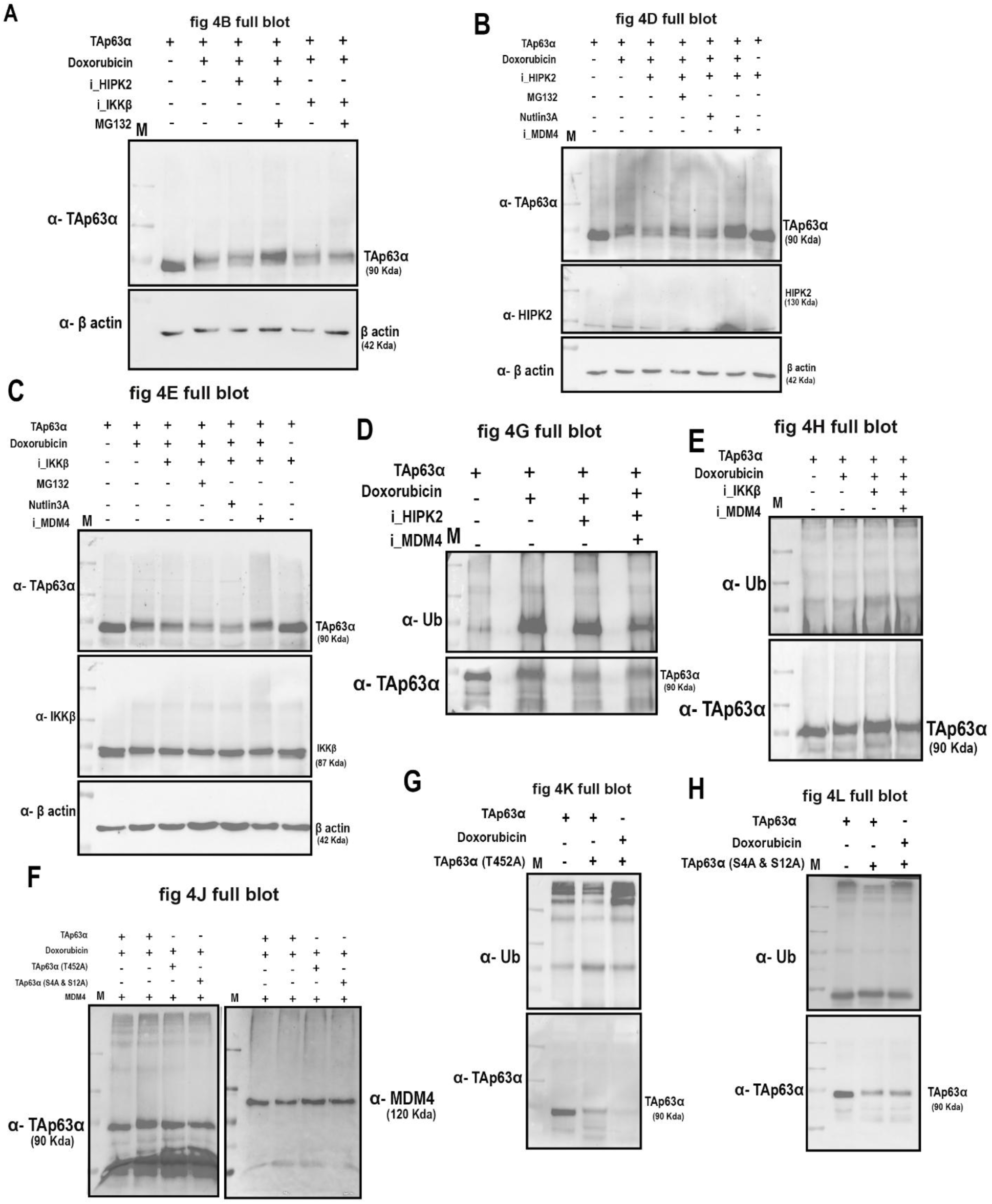

**Table 2:**
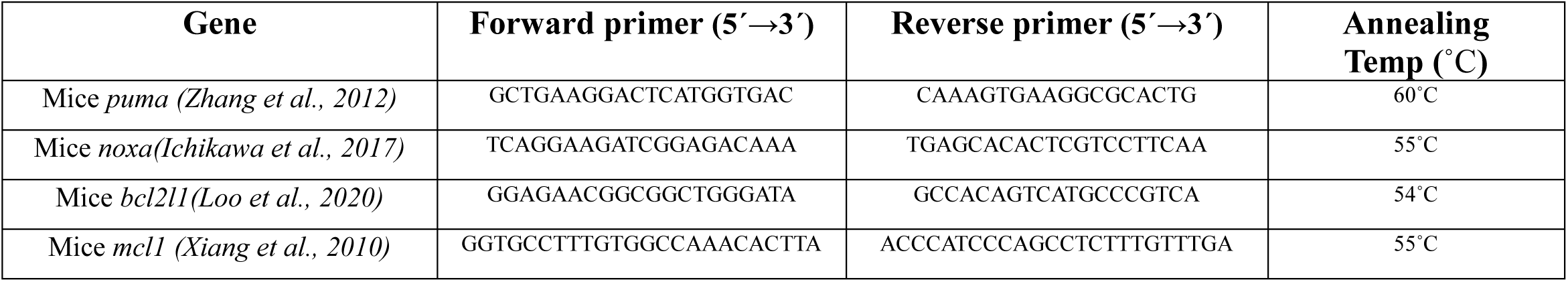
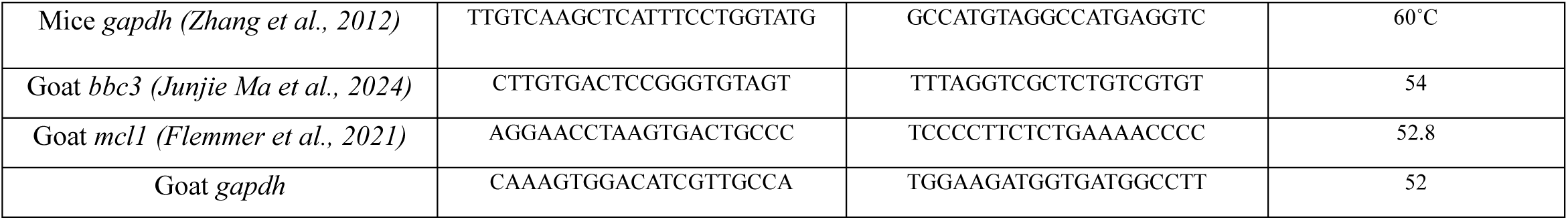
Primer list to amplify the pro and anti-apoptotic genes.

**Table 3:** Inhibitors list and concentration.

| Compound Name | Catalog No. | Company | Final Concentration | Function |
| --- | --- | --- | --- | --- |
| BML-277 | 17552 | Cayman | 10 μM | Chk2 inhibitor |
| PF-670462 | 14588 | Cayman | 1 μM | CK1 inhibitor |
| Protein kinase inhibitors 1 | HY-U00439A | MedChem | 10 μM | HIPK2 inhibitor |
| IKK16 | HY-13687 | MedChem | 10 μM | IKKβ inhibitor |
| Doxorubicin | 15007 | Cayman | 10 μM | DNA damage inducer |
| NSC-207895 | HY-14714 | MedChem | 30 μM | MDM4 inhibitor |
| Nutlin-3 | 10009816 | Cayman | 30 μM | MDM2 inhibitor |
| MG-132 | M3244 | TCI | 30 μM | Proteasome inhibitor |

**Table 4:**
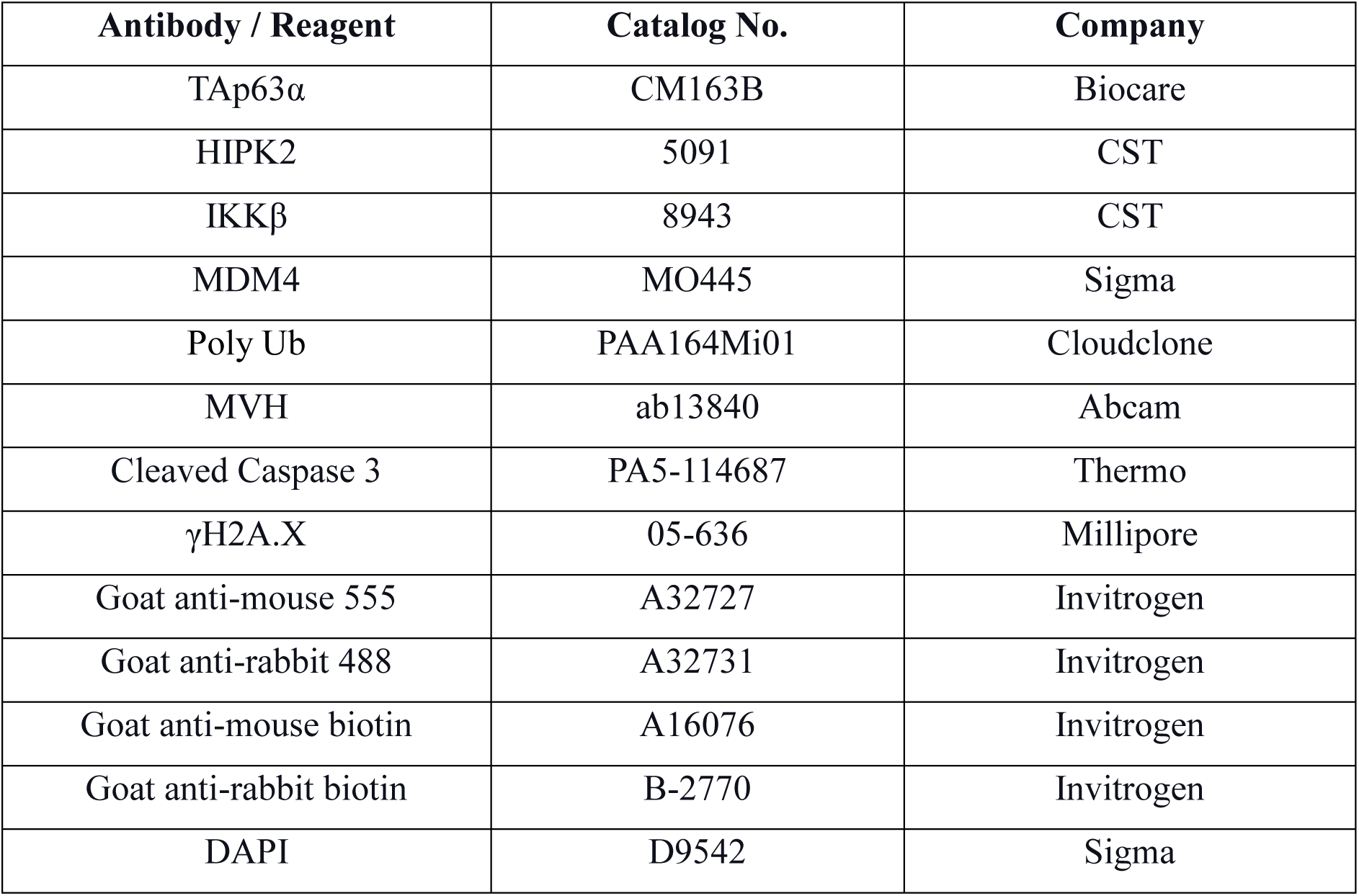
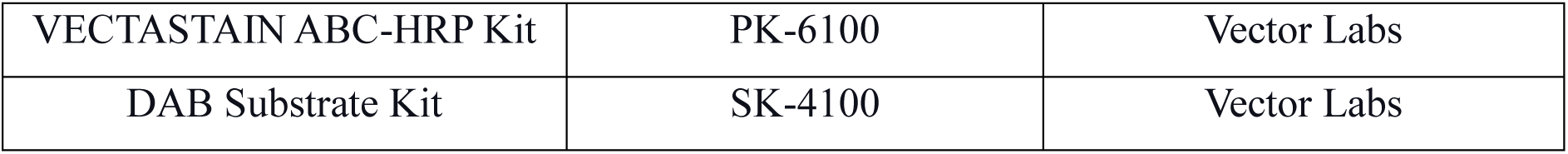
Antibody list.

| Antibody / Reagent | Catalog No. | Company |
| --- | --- | --- |
| TAp63α | CM163B | Biocare |
| HIPK2 | 5091 | CST |
| IKKβ | 8943 | CST |
| MDM4 | MO445 | Sigma |
| Poly Ub | PAA164Mi01 | Cloudclone |
| MVH | ab13840 | Abcam |
| Cleaved Caspase 3 | PA5-114687 | Thermo |
| γH2A.X | 05-636 | Millipore |
| Goat anti-mouse 555 | A32727 | Invitrogen |
| Goat anti-rabbit 488 | A32731 | Invitrogen |
| Goat anti-mouse biotin | A16076 | Invitrogen |
| Goat anti-rabbit biotin | B-2770 | Invitrogen |
| DAPI | D9542 | Sigma |
| VECTASTAIN ABC-HRP Kit | PK-6100 | Vector Labs |
| DAB Substrate Kit | SK-4100 | Vector Labs |

## References

1. Candi, E., Agostini, M., Melino, G., and Bernassola, F. (2014). How the 53 Family Proteins 63 and 73 Contribute to Tumorigenesis: Regulators and Effectors. Human Mutation 35, 702–714. 10.1002/humu.22523.

2. Carroll, J., and Marangos, P. (2013). The DNA damage response in mammalian oocytes. Front Genet 4, 117. 10.3389/fgene.2013.00117.

3. Chillemi, G., Kehrloesser, S., Bernassola, F., Desideri, A., Dötsch, V., Levine, A.J., and Melino, G. (2017). Structural Evolution and Dynamics of the p53 Proteins. Cold Spring Harb Perspect Med 7. 10.1101/cshperspect.a028308.

4. Collins, J.K., and Jones, K.T. (2016). DNA damage responses in mammalian oocytes. Reproduction 152, R15–22. 10.1530/rep-16-0069.

5. Deutsch, G.B., Zielonka, E.M., Coutandin, D., Weber, T.A., Schäfer, B., Hannewald, J., Luh, L.M., Durst, F.G., Ibrahim, M., Hoffmann, J., et al. (2011). DNA damage in oocytes induces a switch of the quality control factor TAp63α from dimer to tetramer. Cell 144, 566–576. 10.1016/j.cell.2011.01.013.

6. Emori, C., Boucher, Z., and Bolcun-Filas, E. (2023). CHEK2 signaling is the key regulator of oocyte survival after chemotherapy. Science Advances 9, eadg0898. doi:10.1126/sciadv.adg0898.

7. Fisher, M.L., Balinth, S., and Mills, A.A. (2020). p63-related signaling at a glance. Journal of Cell Science 133. 10.1242/jcs.228015.

8. Gebel, J., Tuppi, M., Krauskopf, K., Coutandin, D., Pitzius, S., Kehrloesser, S., Osterburg, C., and Dötsch, V. (2017). Control mechanisms in germ cells mediated by p53 family proteins. Journal of Cell Science 130, 2663–2671. 10.1242/jcs.204859.

9. Gebel, J., Tuppi, M., Sänger, N., Schumacher, B., and Dötsch, V. (2020). DNA Damaged Induced Cell Death in Oocytes. Molecules 25. 10.3390/molecules25235714.

10. Gonfloni, S. (2010). DNA damage stress response in germ cells: role of c-Abl and clinical implications. Oncogene 29, 6193–6202. 10.1038/onc.2010.410.

11. Gonfloni, S., Di Tella, L., Caldarola, S., Cannata, S.M., Klinger, F.G., Di Bartolomeo, C., Mattei, M., Candi, E., De Felici, M., Melino, G., and Cesareni, G. (2009). Inhibition of the c-Abl–TAp63 pathway protects mouse oocytes from chemotherapy-induced death. Nature Medicine 15, 1179–1185. 10.1038/nm.2033.

12. Ichikawa, N., Alves, M., Pfeiffer, S., Langa, E., Hernández-Santana, Y.E., Suzuki, H., Prehn, J.H., Engel, T., and Henshall, D.C. (2017). Deletion of the BH3-only protein Noxa alters electrographic seizures but does not protect against hippocampal damage after status epilepticus in mice. Cell Death Dis 8, e2556. 10.1038/cddis.2016.301.

13. Lava Kumar, S., Kushawaha, B., Mohanty, A., Kumari, A., Kumar, A., Beniwal, R., Kiran Kumar, P., Athar, M., Krishna Rao, D., and Rao, H. (2024). Glutathione peroxidase (GPX1) - Selenocysteine metabolism preserves the follicular fluid’s (FF) redox homeostasis via IGF-1- NMD cascade in follicular ovarian cysts (FOCs). Biochim Biophys Acta Mol Basis Dis 1870, 167235. 10.1016/j.bbadis.2024.167235.

14. Lazzari, C., Prodosmo, A., Siepi, F., Rinaldo, C., Galli, F., Gentileschi, M., Bartolazzi, A., Costanzo, A., Sacchi, A., Guerrini, L., and Soddu, S. (2011). HIPK2 phosphorylates ΔNp63α and promotes its degradation in response to DNA damage. Oncogene 30, 4802–4813. 10.1038/onc.2011.182.

15. Liao, J.M., Zhang, Y., Liao, W., Zeng, S.X., Su, X., Flores, E.R., and Lu, H. (2013). IκB kinase β (IKKβ) inhibits p63 isoform γ (TAp63γ) transcriptional activity. J Biol Chem 288, 18184–18193. 10.1074/jbc.M113.466540.

16. Livera, G., Petre-Lazar, B., Guerquin, M.-J., Trautmann, E., Coffigny, H., and Habert, R. (2008). p63 null mutation protects mouse oocytes from radio-induced apoptosis. REPRODUCTION 135, 3–12. 10.1530/rep-07-0054.

17. Loo, L.S.W., Soetedjo, A.A.P., Lau, H.H., Ng, N.H.J., Ghosh, S., Nguyen, L., Krishnan, V.G., Choi, H., Roca, X., Hoon, S., and Teo, A.K.K. (2020). BCL-xL/BCL2L1 is a critical anti-apoptotic protein that promotes the survival of differentiating pancreatic cells from human pluripotent stem cells. Cell Death & Disease 11, 378. 10.1038/s41419-020-2589-7.

18. Luan, Y., Xu, P., Yu, S.Y., and Kim, S.Y. (2021). The Role of Mutant p63 in Female Fertility. Int J Mol Sci 22. 10.3390/ijms22168968.

19. Luan, Y., Yu, S.Y., Abazarikia, A., Dong, R., and Kim, S.Y. (2022). TAp63 determines the fate of oocytes against DNA damage. Sci Adv 8, eade1846. 10.1126/sciadv.ade1846.

20. MacPartlin, M., Zeng, S.X., and Lu, H. (2008). Phosphorylation and stabilization of TAp63gamma by IkappaB kinase-beta. J Biol Chem 283, 15754–15761. 10.1074/jbc.M801394200.

21. Mahajan, N. (2015). Fertility preservation in female cancer patients: An overview. J Hum Reprod Sci 8, 3–13. 10.4103/0974-1208.153119.

22. Marcel, V., Dichtel-Danjoy, M.L., Sagne, C., Hafsi, H., Ma, D., Ortiz-Cuaran, S., Olivier, M., Hall, J., Mollereau, B., Hainaut, P., and Bourdon, J.C. (2011). Biological functions of p53 isoforms through evolution: lessons from animal and cellular models. Cell Death & Differentiation 18, 1815–1824. 10.1038/cdd.2011.120.

23. Mohanty, A., Kumari, A., Kumar. S. L., Kumar, A., Birajdar, P., Beniwal, R., Athar, M., Kumar P, K., and Prasada Rao, H.B.D. (2024). Cathepsin B regulates ovarian reserve quality and quantity via mitophagy by modulating IGF1R turnover. bioRxiv, 2024.2002.2014.580410. 10.1101/2024.02.14.580410.

24. Moll, U.M., and Slade, N. (2004). p63 and p73: roles in development and tumor formation. Mol Cancer Res 2, 371–386.

25. Osterburg, C., and Dötsch, V. (2022). Structural diversity of p63 and p73 isoforms. Cell Death Differ 29, 921–937. 10.1038/s41418-022-00975-4.

26. Petitjean, A., Ruptier, C., Tribollet, V., Hautefeuille, A., Chardon, F., Cavard, C., Puisieux, A., Hainaut, P., and Caron de Fromentel, C. (2007). Properties of the six isoforms of p63: p53-like regulation in response to genotoxic stress and cross talk with ΔNp73. Carcinogenesis 29, 273–281. 10.1093/carcin/bgm258.

27. Rinaldi, V.D., Bloom, J.C., and Schimenti, J.C. (2020). Oocyte Elimination Through DNA Damage Signaling from CHK1/CHK2 to p53 and p63. Genetics 215, 373–378. 10.1534/genetics.120.303182.

28. Singh, A.K., Kumar, S.L., Beniwal, R., Mohanty, A., Kushwaha, B., and Prasada Rao, H.B.D. (2023). Influence of the Ovarian Reserve and Oocyte Quality on Livestock Fertility. In Sustainable Agriculture Reviews 59: Animal Biotechnology for Livestock Production 3, V.K. Yata, A.K. Mohanty, and E. Lichtfouse, eds. (Springer Nature Switzerland), pp. 201–240. 10.1007/978-3-031-21630-5_4.

29. Suh, E.-K., Yang, A., Kettenbach, A., Bamberger, C., Michaelis, A.H., Zhu, Z., Elvin, J.A., Bronson, R.T., Crum, C.P., and McKeon, F. (2006). p63 protects the female germ line during meiotic arrest. Nature 444, 624–628. 10.1038/nature05337.

30. Tuppi, M., Kehrloesser, S., Coutandin, D.W., Rossi, V., Luh, L.M., Strubel, A., Hötte, K., Hoffmeister, M., Schäfer, B., De Oliveira, T., et al. (2018). Oocyte DNA damage quality control requires consecutive interplay of CHK2 and CK1 to activate p63. Nature Structural & Molecular Biology 25, 261–269. 10.1038/s41594-018-0035-7.

31. Xiang, Z., Luo, H., Payton, J.E., Cain, J., Ley, T.J., Opferman, J.T., and Tomasson, M.H. (2010). Mcl1 haploinsufficiency protects mice from Myc-induced acute myeloid leukemia. J Clin Invest 120, 2109–2118. 10.1172/JCI39964.

32. Zhang, F., Li, Y., Tang, Z., Kumar, A., Lee, C., Zhang, L., Zhu, C., Klotzsche-von Ameln, A., Wang, B., Gao, Z., et al. (2012). Proliferative and Survival Effects of PUMA Promote Angiogenesis. Cell Reports 2, 1272–1285. 10.1016/j.celrep.2012.09.023.

